# Multifunctional requirements for ERK1/2 signaling in the development of ganglionic eminence derived glia and cortical inhibitory neurons

**DOI:** 10.1101/2022.08.02.502073

**Authors:** Sara J. Knowles, Michael C. Holter, Guohui Li, George R. Bjorklund, Katherina P. Rees, Johan S. Martinez-Fuentes, Kenji J. Nishimura, Ariana E. Afshari, Noah Fry, April M Stafford, Daniel Vogt, Marco Mangone, Trent Anderson, Jason M. Newbern

**Author notes:** Corresponding Author address: School of Life Sciences, PO Box 84501 Arizona State University Tempe, AZ 85287-4501, USA.

## Abstract

The RAS/RAF/MEK/ERK1/2 intracellular signaling pathway is activated by numerous cues during brain development and dysregulated in neurodevelopmental syndromes, particularly the RASopathies and certain forms of autism. Cortical excitatory/inhibitory imbalance is thought to be critical in the neuropathogenesis of these conditions. However, the developmental functions of ERK1/2 signaling in cortical inhibitory neurons (CINs) and other medial ganglionic eminence (MGE)-derived non-neuronal cells are poorly understood. Here, we genetically modulated ERK1/2 signaling in mouse MGE neural progenitors or GABAergic neurons in vivo. We find that MEK-ERK1/2 signaling is essential for regulating MGE-derived oligodendrocyte number in the anterior commissure. While *Erk1/2* inactivation does not alter CIN number, we discovered a significant and persistent reduction in somatostatin, but not parvalbumin, expression in a subset of CINs. ERK1/2 signaling is also necessary for chemogenetic activity-dependent FOSB expression in CINs in vivo. Interestingly, one week of chronic chemogenetic stimulation in juvenile or adult animals partially rescues the decrease in somatostatin expression in *Erk1/2* mutant CINs. Our data demonstrate ERK1/2 signaling is required for the establishment of MGE-derived glia, whereas in CINs, ERK1/2 drives activity dependent-responses and the expression of somatostatin in a subset of neurons.

## Introduction

The canonical RAS/RAF/MEK/ERK1/2 intracellular kinase cascade is a ubiquitous pathway activated by a wide array of extracellular signals (Lavoie et al., 2020). Mutations in Receptor Tyrosine Kinase (RTK)-associated effectors and upstream regulators of Extracellular Regulated Kinase (ERK1/2, aka *MAPK3/1*) signaling cause a family of developmental syndromes collectively termed the “RASopathies” (Rauen, 2013). Individuals with RASopathies frequently exhibit craniofacial abnormalities, cardiac malformations, and increased cancer risk, in addition to neurological conditions such as epilepsy, developmental delay, intellectual disability, and autism (Tidyman & Rauen, 2016). Most RASopathy mutations increase ERK1/2 signaling, however, microdeletions that disrupt *Erk1* or *Erk2* expression have been linked to neurocognitive delay, intellectual disability, and autism (Ben-Shachar et al., 2008; Fernandez et al., 2010; Linhares et al., 2016; Newbern et al., 2008; Nowaczyk et al., 2014; Pucilowska et al., 2015; Rauen, 2013; Saitta et al., 2004; Samuels et al., 2009; Sánchez et al., 2020). Moreover, ERK1/2 signaling is a point of functional convergence for several autism and schizophrenia-associated copy number variants (Blizinsky et al., 2016; Courchesne et al., 2020; Heavner & Smith, 2020; Lord et al., 2020; Moyses-Oliveira et al., 2020; Rosina et al., 2019). Understanding the normal functions of ERK1/2 signaling in the developing forebrain may be instructive for defining neuropathological mechanisms in multiple neurodevelopmental syndromes.

The patterning of the forebrain is heavily dependent upon canonical trophic cues known to activate RTK signaling, but the precise functions of downstream kinase cascades are less clear (Assimacopoulos et al., 2003; Faedo et al., 2010; Maisonpierre et al., 1990; O’Leary et al., 2007). Studies of the mouse dorsal pallium have shown that FGF and ERK1/2 signaling via ETV5 are crucial for cortical gliogenesis, while neurogenesis is relatively less affected (Dinh Duong et al., 2019; Fyffe-Maricich et al., 2011; S. Li et al., 2014; X. Li et al., 2012; Pucilowska et al., 2012). Interestingly, cell-specific *Mek1/2*-deletion in embryonic cortical excitatory neurons leads to subtype-specific deficits in morphological and electrophysiological properties during the early postnatal period (L. Xing et al., 2016). In contrast to mouse excitatory neurons, GABAergic cortical inhibitory neurons (CINs) are primarily derived from progenitor domains in the medial and caudal ganglionic eminences (MGE and CGE) (Wonders & Anderson, 2006). CIN patterning cues, migratory paths, master transcription factors, and expression levels of ERK1/2 pathway components differ significantly from dorsally derived glutamatergic neurons (Mardinly et al., 2016; Mayer et al., 2018; Ryu, Kang, et al., 2019; Ryu, Kim, et al., 2019; Talley et al., 2021). It is unknown whether ERK1/2 signaling has similar or unique functions in the development of glial and neuronal derivatives of the ganglionic eminences.

CINs exhibit substantial heterogeneity in their morphology, gene expression profile, electrophysiological properties, and contributions to cognition (Bomkamp et al., 2019; DeFelipe et al., 2013; Gour et al., 2021; Lim et al., 2018; Mayer et al., 2018; Petros et al., 2015). The two largest classes of CINs are identified by parvalbumin (PV) or somatostatin (SST) expression and are distinguished by unique electrophysiological properties (Dehorter et al., 2015; Lim et al., 2018; Okaty et al., 2009; Pan et al., 2018; Urban-Ciecko & Barth, 2016). Differences in their generation and local morphogen gradients bias progeny towards specific subtypes (McKenzie et al., 2019; Miyoshi et al., 2007; Petros et al., 2015). Trophic cues and neurotransmitters (e.g. BDNF/TrkB, Neuregulin-1/ErbB4, glutamate/NMDAR) are also important extracellular influences on CIN development (Cancedda et al., 2007; De Marco García et al., 2011; Exposito-Alonso et al., 2020; Fazzari et al., 2010; Glorioso et al., 2006; Hong et al., 2008; Huang et al., 1999; Itami et al., 2007; Miller et al., 2011). While the transcriptional mechanisms of CIN development have been intensely studied, the signaling pathways that serve as intermediaries between extracellular stimulation and gene expression changes are relatively less clear. Recent work has shown that hyperactive MEK-ERK1/2 and PI3K-linked signaling pathways have critical roles in CIN development and subtype specification (Angara et al., 2020; Knowles et al., 2022; Malik et al., 2019; Vogt et al., 2015). However, the field still lacks crucial information on whether ERK1/2 signaling is necessary for CIN subtype identity.

The initial stages of postnatal development are characterized by a series of activity-dependent events in which CINs play a critical role (Ben-Ari et al., 2007; Cancedda et al., 2007; Close et al., 2012; Denaxa et al., 2012). Early spontaneous activity causes the release of GABA, which functions as a neurotransmitter and trophic cue to depolarize neurons and promote connections in the nascent cortical circuit (Ben-Ari et al., 2007; Owens & Kriegstein, 2002). Later stages of patterned activity are especially important for establishing the timing of critical periods and excitatory/inhibitory tone in cortical networks (Reh et al., 2020). Activity-dependent responses have been well studied in cortical excitatory neurons, where glutamatergic activation of ERK1/2 has been shown to modulate synaptic strength (Cristo et al., 2001; Thomas & Huganir, 2004; Wu et al., 2001; J. Xing et al., 1996; Yap & Greenberg, 2018). CINs express different levels of glutamate receptor subunits, calcium-binding proteins, and calcium responsive kinases that modulate intracellular signaling responses to glutamate (Cohen et al., 2016; Matthews et al., 1994; Paul et al., 2017; Sik et al., 1998; Tasic et al., 2016; Topolnik, 2012). Indeed, many features of the activity-dependent transcriptome in CINs differ from excitatory neurons (Guenthner et al., 2013; Hrvatin et al., 2018; Mardinly et al., 2016; Spiegel et al., 2014). Nonetheless, the phosphorylation-dependent signaling requirements for ERK1/2 in CINs are difficult to assess with transcriptomic approaches.

Studying the functions of ERK1/2 specifically in ventral forebrain derivatives using germline knockouts of *Erk1* and *Erk2* is complicated by a combination of embryonic lethality, compensatory interactions, and non-cell autonomous effects (Newbern et al., 2011; Samuels et al., 2008; Selcher, 2001). Here, we used cell-specific genetic modifications to conditionally inactivate ERK1/2 signaling in medial ganglionic eminence (MGE) progenitors (*Nkx2.1^Cre^*) and GABAergic neurons (*VGAT^Cre^*) in embryonic mice. We show that ERK1/2 signaling is a critical modulator of MGE-derived oligodendrocyte number in juvenile mice. In contrast, *Erk1/2* appears dispensable for early tangential migration of CINs, the expression of core cortical GABAergic neuron genes, and global CIN number. Further analysis of postnatal stages of cortical maturation, however, revealed that the expression of SST protein in a subset of CINs requires ERK1/2 signaling. We also found that *Erk1/2* is vital for activity-dependent FOSB expression in CINs following chemogenetic stimulation in vivo. Moreover, chronically increasing CIN activity during early postnatal stages or in adulthood was sufficient to partially rescue the proportion of CINs that express SST. Our data reveal previously undescribed roles for ERK1/2 signaling in select aspects of ventral forebrain glial development and the maturation of CINs. These results may also inform our understanding of the cellular events underlying neurodevelopmental disorders linked to aberrant ERK1/2 signaling.

## Results

### ERK1/2 signaling regulates the number of ventral forebrain-derived oligodendrocytes

The ERK1/2 signaling pathway is an established regulator of dorsal pallium derived glial number, but its role in glial specification from subpallial progenitor domains remains less clear (Dinh Duong et al., 2019; X. Li et al., 2012). Neural progenitors within the subpallial medial ganglionic eminence (MGE) produce both neurons and glia after regional patterning by the transcription factor, Nkx2.1, at ∼E8.5 (Shimamura & Rubenstein, 1997). *Nkx2.1^Cre^*mice provide a valuable tool to selectively induce genetic modifications in MGE-derived glia and cortical inhibitory neurons (CINs) early in development. To completely inhibit ERK1/2 signaling and fluorescently label MGE derivatives, we generated quadruple genetically modified mice with germline *Erk1*/*Mapk3* deletion (*Erk1*^-/-^), loxp-flanked ("floxed”) *Erk2/Mapk1* exon 2 alleles (*Erk2^fl/fl^*), *Nkx2.1^Cre^ ^+/-^*, and a Cre-dependent tdTomato/RFP reporter, *Ai9^+/-^* (see *Genetically modified mice* in Materials and Methods) (Madisen et al., 2010; Newbern et al., 2008; Samuels et al., 2008; Selcher, 2001; L. Xing et al., 2016; Xu et al., 2008). *Nkx2.1^Cre^*; *Ai9* mice showed an overall pattern of RFP-labeled neurons and glia consistent with previous reports at P14 (**Figure 1A**) and E13.5 (**Figure 1-figure supplement 1A**) (Madisen et al., 2010; Xu et al., 2008). Mutant mice with complete *Nkx2.1^Cre^*-mediated *Erk1/2* inactivation (*Erk1^-/-^; Erk2^fl/fl^; Nkx2.1^Cre^; Ai9*) were born at normal Mendelian ratios, survived into adulthood, and exhibited a reduction in ERK2 protein in CINs compared with *Erk1^+/+^; Erk2^wt/wt^; Nkx2.1^Cre^*; *Ai9* (WT) or *Erk1^-/-^; Erk2^fl/wt^; Nkx2.1^Cre^; Ai9* (HET control) mice of the same age (**Figure 1-figure supplement 1B-J**). Nearby cells, presumptive excitatory neurons, often showed higher levels of ERK2 protein expression, consistent with previous reports (Holter et al., 2021; Mardinly et al., 2016). Past research has demonstrated that DRG sensory or cortical excitatory neuron directed *Erk1^-/-^; Erk2^fl/wt^; Cre* expressing mice were relatively normal (Newbern et al., 2011; L. Xing et al., 2016). *Erk1^-/-^; Erk2^fl/wt^; Nkx2.1^Cre^; Ai9* mice were also grossly intact and used as littermate controls for certain experiments.

**Figure 1.**
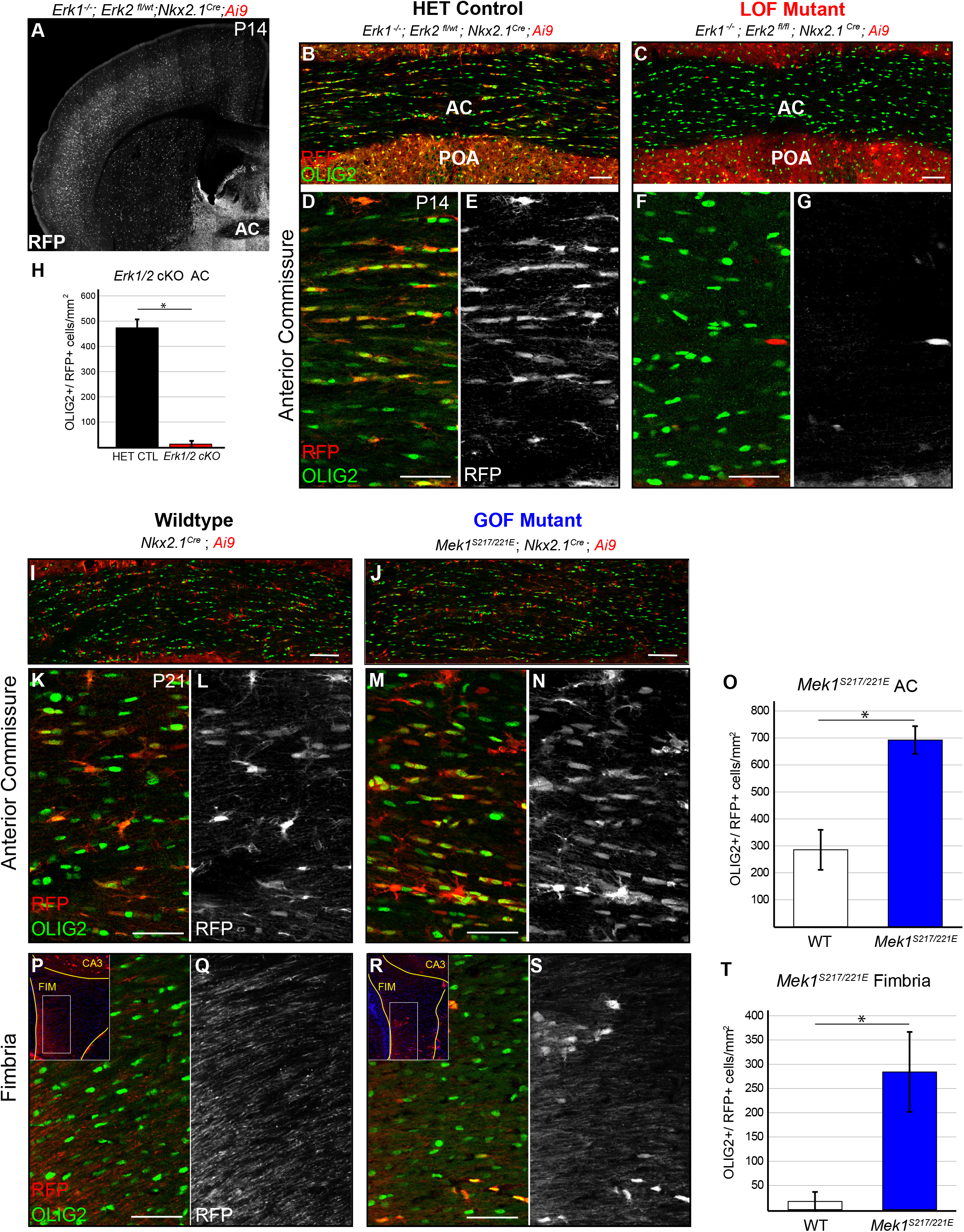
ERK1/2 signaling regulates MGE-derived oligodendrocyte number. **(A)** Anatomical image of a P14 *Erk1^-/-^; Erk2^fl/wt^; Nkx2.1^Cre^*; *Ai9^+/-^*coronal section showing the expected neuroanatomical pattern of *Nkx2.1^Cre^*-mediated recombination. Note the presence of *Nkx2.1^Cre^*-derived oligodendrocytes in the anterior commissure at this stage. **(B-H)** Representative confocal images of the anterior commissure at P14 display a significant decrease in the density of OLIG2^+^/RFP^+^ co-labeled cells in *Erk1^-/-^; Erk2^fl/fl^; Nkx2.1^Cre^; Ai9* mutants (C, F-G) compared to HET controls (B, D-E) (quantification in **H**; N=3, mean ± SEM, *student’s t-test, *p*=.0002) (scale bars 100µm (B-C), 50µm (D-G)). **(I-O)** Confocal images of the anterior commissure at P21 revealed a significant increase in the number of OLIG2^+^/RFP^+^ cells in *Mek1^S217/221E^; Nkx2.1^Cre^; Ai9* mutants (J, M-N) relative to wildtype (I, K-L) (scale bar = 100 µm I-J, 50 µm K-N) (Quantification in **O**; N=3-4, mean ± SEM, *student’s t-test, *p*=.004). **(P-T)** We observed ectopic expression of OLIG2^+^/RFP^+^ cells in the fimbria of *Mek1^S217/221E^; Nkx2.1^Cre^; Ai9* mutants (scale bar = 50 µm) (quantification in **T**; N=3-4, mean ± SEM, *student’s t-test, *p*=.005).

The anterior commissure is a ventral white matter tract populated by glia from multiple progenitor domains including the MGE (Kessaris et al., 2006; Minocha et al., 2015). Glia from *Erk1^-/-^; Erk2^fl/fl^; Nkx2.1^Cre^; Ai9* mutants had less ERK2 immunoreactivity than *Erk1^-/-^; Erk2^fl/wt^; Nkx2.1^Cre^; Ai9* HET controls (**Figure 1-figure supplement 2B-G**). We used a marker of the oligodendrocyte lineage, OLIG2, to quantify the density of oligodendrocytes and OPCs at postnatal day 14 (P14). We found a significant reduction in OLIG2^+^/RFP^+^ co-expressing cells in the anterior commissure of *Erk1^-/-^; Erk2^fl/fl^; Nkx2.1^Cre^; Ai9* mutants when compared with *Erk1^-/-^; Erk2^fl/wt^; Nkx2.1^Cre^; Ai9* littermate controls (CTL 473 +/- 34 cells/mm^2^, MUT 13 +/- 13 cells/mm^2^, *p*=.0002, student’s t-test) (**Figure 1 B-H, Figure 1-figure supplement 2A**). GFAP^+^/RFP^+^ astrocyte density was similarly reduced (CTL 84 +/- 9 cells/mm^2^, MUT 19 +/- 9 cells/mm^2^, *p*=.016, student’s t-test) (**Figure 1-figure supplement 2H-J**), suggesting that ERK1/2 signaling is necessary for MGE-derived astrocyte and oligodendrocyte number.

RASopathies are primarily caused by mutations that hyperactivate ERK1/2, and often other, signaling cascades. To determine if hyperactive MEK-ERK1/2 signaling is sufficient to modulate MGE-derived glial density in the ventral forebrain, we generated mice expressing a Cre-dependent hyperactivating *Mek1^S217/221E^* mutation (*CAG-loxpSTOPloxp-Mek1^S217/221E^)*. *Mek1^S217/221E^; Nkx2.1^Cre^; Ai9* mutants display relatively increased MEK1 levels in RFP^+^ CINs and glia (**Figure 1-figure supplement 1K-P, Figure 1-figure supplement 2K-P**) (Alessi et al., 1994; Bueno, 2000; Cowley, 1994; Holter et al., 2021; Klesse et al., 1999; Krenz et al., 2008; Lajiness et al., 2014; X. Li et al., 2012). In contrast to *Erk1/2* inactivation, *Mek1^S217/221E^; Nkx2.1^Cre^; Ai9* mutants showed an increase in OLIG2^+^/RFP^+^ cell density in the anterior commissure at P21 compared with *Nkx2.1^Cre^; Ai9* littermate controls (WT 285 +/- 74 cells/mm^2^, *Mek1^S217/221E^* 692 +/- 51 cells/mm^2^, *p*=.004, student’s t-test) (**Figure 1I-O**, **Figure 1-figure supplement 2A**). In addition, we observed ectopic MGE-derived glial profiles in the fimbria of *Mek1^S217/221E^; Nkx2.1^Cre^; Ai9* mutants (**Figure 1P-T**). However, we did not observe an increase in RFP+/GFAP+ cell density in the anterior commissure of these animals (**Figure 1-figure supplement 2Q-S**). These data indicate that increased MEK-ERK1/2 signaling enhances MGE-derived oligodendrocyte number.

### Reduced ERK1/2 signaling does not alter total CIN number in the somatosensory cortex

ERK1/2 signaling is necessary for the survival and physiological maturation of select excitatory neuron subpopulations, but its requirement for CIN development is unclear (L. Xing et al., 2016). In the primary somatosensory cortex of *Erk1^-/-^; Erk2^fl/fl^; Nkx2.1^Cre^; Ai9* mice, we found no difference in the density of *Nkx2.1^Cre^*-recombined cells compared with HET controls at P14 (**Figure 2A-E**, *p*=0.476, student’s t-test) and P60 (*p*=0.288, student’s t-test, data not shown). *Nkx2.1^Cre^* does not recombine in the fraction of CINs derived from the lateral or caudal ganglionic eminences (Kessaris et al., 2006; Talley et al., 2021). These data suggest ERK1/2 is not required for the generation of the subset of CINs derived from the MGE.

**Figure 2.**
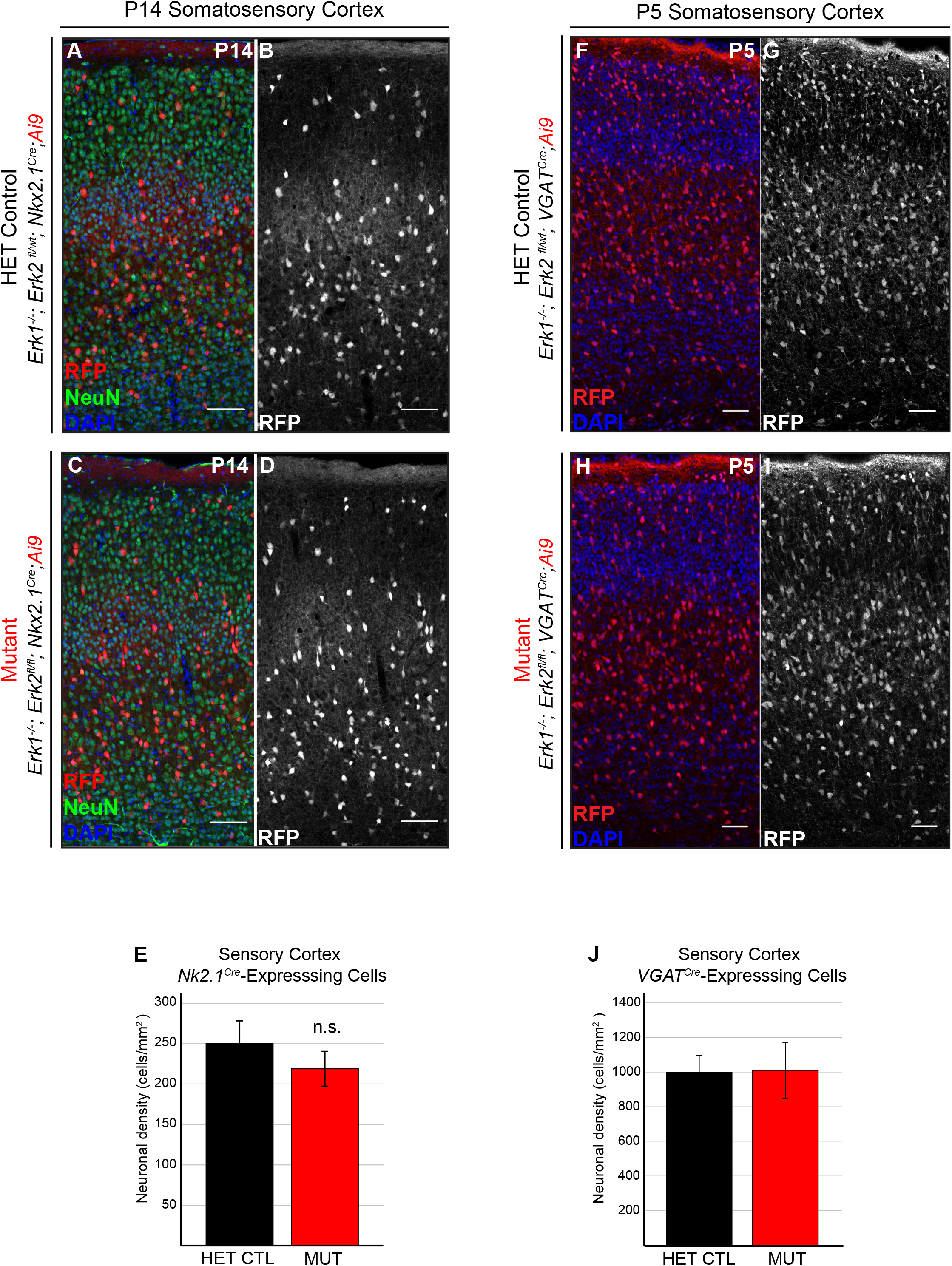
*Erk1/2* deletion does not alter CIN number. **(A-E)** Representative images of P14 somatosensory cortex in HET control (A-B) and *Erk1^-/-^; Erk2^fl/fl^; Nkx2.1^Cre^; Ai9* (C-D) mutant animals, showing similar density of *Nkx2.1^Cre^*-positive cells (quantification in **E**, N=4, mean +/- SEM, student’s t-test, *p*=0.476) (scale bar = 50µm). **(F-J)** At P5, heterozygote control (F-G) and *Erk1^-/-^; Erk2^fl/fl^; VGAT^Cre^; Ai9* mutant animals (H-I) showed no significant difference in the density of *VGAT^Cre^*-positive cells in the somatosensory cortex (quantification in **J**, N=4, mean +/- SEM, student’s t-test *p*=.955) (scale bar = 50µm).

To comprehensively assess the role of *Erk1/2* in all postmitotic GABAergic neurons, we generated conditional mutants using *VGAT/Slc32a1^Cre^*(Madisen et al., 2010; Vong et al., 2011). *VGAT^Cre^* is expressed in all GABAergic neurons early in development. We found that *Erk1^-/-^; Erk2^fl/fl^; VGAT^Cre^; Ai9* mutant mice were born at normal Mendelian ratios but exhibited profound growth delay by the end of the first postnatal week and lethality in the second week (N=10). No significant difference in the density of *VGAT^Cre^*-recombined cells was observed between P5 *Erk1^-/-^; Erk2^fl/fl^; VGAT^Cre^; Ai9* mutant and *Erk1^-/-^; Erk2^fl/wt^; VGAT^Cre^; Ai9* HET control cortices (CTL 998 +/- 98 cells/mm^2^, MUT 1010 +/- 162 cells/mm^2^, *p*=.955, student’s t-test) (**Figure 2F-J**). Together, these data suggest that ERK1/2 signaling is dispensable for the initial establishment and maintenance of CIN number.

### ERK1/2 deletion alters SST expression in a subset of inhibitory neurons

CINs are a heterogeneous population of neurons that can be divided into numerous subtypes with overlapping but distinct transcriptional profiles. Although CIN number was unaltered in the cortex following ERK1/2 deletion, gene expression changes in these cells remained unknown. To study the CIN translatome, we employed a Cre-dependent ‘Ribotag’ mouse that expresses an HA-fused ribosomal protein, RPL22^HA^, for translating ribosome affinity purification followed by RNA sequencing (Sanz et al 2009). Whole cortices were dissected from P7.5 *Erk1^-/-^; Erk2^fl/fl^; VGAT^Cre^; Rpl22^HA^* (MUT), *Erk1^-/-^; Erk2^fl/wt^; VGAT^Cre^; Rpl22^HA^*(HET), or wildtype controls (*Erk1^+/+^; Erk2^wt/wt^; VGAT^Cre^; Rpl22^HA^*) (WT). Following homogenization, HA-containing ribosomes were immunoprecipitated and RNA was purified from cortical (INPUT) or immunoprecipitated (IP) samples, amplified, and sequenced. As expected, we observed a near complete absence of reads complementary to *Erk1/Mapk3* exons 1-6 in *Erk1^-/-^* animals and a gene dose dependent reduction of *Erk2/Mapk1* exon 2 in IP fractions (**Figure 3-figure supplement 1A**). A comparison of the >3400 differentially expressed genes between IP and INPUT fractions within each genotype revealed that IP fractions exhibited significant enrichment of *Cre* and well-established core GABAergic genes (*Gad1/2*, *Slc32a1*, *Erbb4*, *Dlx1/2/5/6*), as well as significant depletion of genes expressed in glia (*Fabp7, Sox10, Pdgfra, S100b*) and excitatory neurons (*Neurod1/6, Camk2a*) (**Table S1** and **Figure 3-figure supplement 1B-C**) (Hrvatin et al., 2018; Mardinly et al., 2016; Mi et al., 2018; Paul et al., 2017; Tasic et al., 2016). These data indicate the Ribotag approach significantly enriched samples for CIN-derived mRNA.

We next examined the differential expression of genes between control (HET) and mutant (MUT) IP, CIN-enriched samples, particularly those important for GABAergic specification and subtype identity (**Table S2**). A significant decrease in known ERK1/2 target genes *Spry4*, *Etv5*, and *Egr2* was noted in MUT IP samples (**Figure 3A-magenta**) (X. Li et al., 2012; Newbern et al., 2011; L. Xing et al., 2016). However, the expression of master transcriptional regulators of global CIN identity (*Lhx6*, *Dlx2/5/6)* and GABAergic metabolism *(Gad1/2*, *Slc32a1*) was not significantly different between HET and MUT IP samples (**Figure 3A-green**). GO analysis detected a significant overrepresentation of differentially expressed genes in the category of “regulation of cell differentiation” (GO:0045595, adj. *p* =.002) and “nervous system development” (GO:0030182, adj. *p* =.04). Indeed, we observed changes in the expression of genes linked to neural maturation (*Notch1, Smo, Fgf18, Bdnf*) and CIN subtype specification (*Mafb, Sst, Sp9, Crhbp, Grin2d*). Together, these data indicate that ERK1/2 inactivation does not significantly disrupt core mediators of CIN identity but may have selective effects on later stages of subtype maturation.

**Figure 3.**
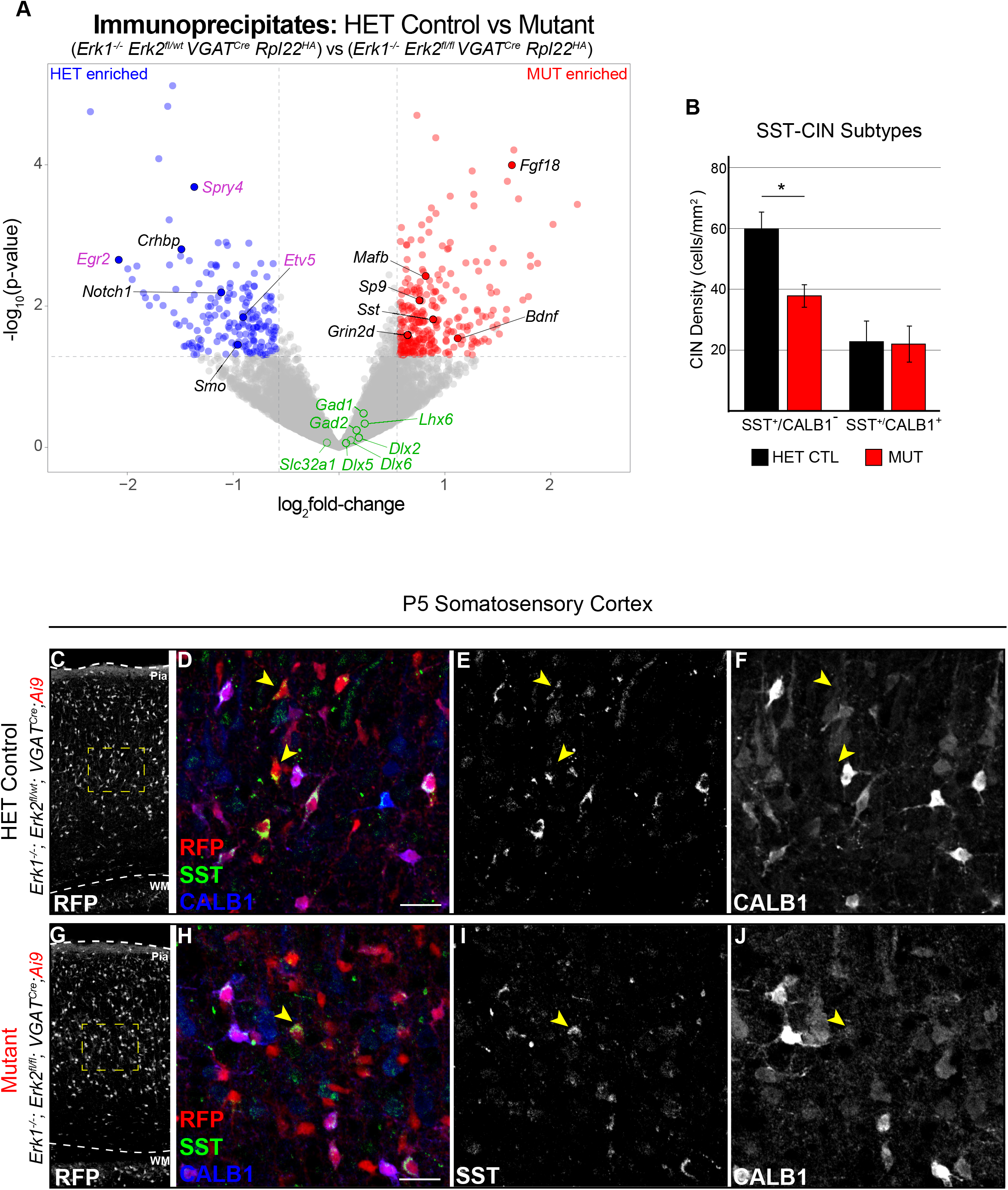
Loss of ERK1/2 does not alter the initial expression of generic regulators of GABAergic function. **(A)** HET IP vs MUTANT IP volcano plot of expressed protein-coding genes in Ribotag, CIN-enriched samples illustrates the significant reduction in ERK1/2 pathway target genes in *Erk1/2* inactivated (*Erk1^-/-^; Erk2^fl/fl^; VGAT^Cre^; Rpl22^HA^)* immunoprecipitated fractions (magenta - *Spry4, Etv5, Egr2*) (N=3, *p*<.05). A significant difference was not detected in the expression of master regulators of CIN specification (green - *Lhx6, Dlx2/5/6*) and key GABA processing genes (green - *Gad1/2, Slc32a1*). However, the expression CIN-subtype enriched genes (*Crhbp, Sst, Mafb, Grin2d, Sp9*) and canonical developmental signals (*Notch1, Smo, Fgf18, Bdnf*) were significantly altered in mutant CINs. **(B-J)** Immunolabeling analysis of CIN subtype markers revealed a significant reduction in SST^+^/CALB**^-^** cell density (yellow arrowheads) in mutants, while SST^+^/CALB^+^ co-labeled cell density was unchanged (quantification in **B**, N=4, mean ± SEM, *student’s t-test *p*=0.017). Representative images in (D-F, H-J) are from layer 5 primary somatosensory cortex (scale bar = 25µm).

We further analyzed CIN subtypes in *Erk1^-/-^; Erk2^fl/fl^; VGAT^Cre^; Ai9* brains at P5, prior to the observed lethality. At this stage, SST, a canonical marker of a subset of regular spiking (RS) CINs, and Calbindin 1 (CALB1), a calcium binding protein co-expressed in a subset of CINs partially overlapping with SST, are expressed (Forloni et al., 1990; Kawaguchi, 1997; Marsh et al., 2016; Martin et al., 2017; Mi et al., 2018; Munguba et al., 2019; Nigro et al., 2018; Tasic et al., 2016). We assessed these two classes of SST-expressing CINs and found a statistically significant reduction in SST^+^/CALB1**^-^** cells in mutants relative to littermate controls (*p*=0.017, student’s t-test), but no change in SST^+^/CALB1^+^ co-expressing cells (*p*=0.938, student’s t-test) (**Figure 3B-J**). The early lethality of the *Erk1^-/-^; Erk2^fl/fl^; VGAT^Cre^; Ai9* mice prohibited our ability to study fully mature CIN markers, as PV is not expressed until the second postnatal week and only a subset of RS CINs have initiated SST protein expression (del Rio et al., 1994; Forloni et al., 1990; Huang et al., 1999). Nonetheless, our data are consistent with a role for ERK1/2 in the early development of SST-CIN subtypes in the first postnatal week.

The broad recombination pattern of *VGAT^Cre^* throughout the entire nervous system leaves open the possibility of non-cell-autonomous effects on CIN development. We therefore performed more detailed studies of subtype specification in *Nkx2.1^Cre^* mutants, where recombination occurs in a smaller subset of CINs, but survival extends into adulthood. Quantification of PV-CIN number in P14 and adult *Erk1^-/-^; Erk2^fl/fl^; Nkx2.1^Cre^; Ai3* mice revealed no differences in the proportion of *Nkx2.1^Cre^*-derived CINs expressing PV or their somal size (CTL: 181.7 +/- 4.4µm^2^, MUT: 187.4 +/- 5.0µm^2^, mean +/- SEM, N=105 cells/group, student’s t-test, *p*=0.398) (**Figure 4A-C, E-G, I-K, M-O, Q**). Notably, we observed a significant reduction in the proportion of *Nkx2.1^Cre^*-derived, SST-expressing CINs at P14, an effect which persisted into adulthood (Figure 4D, H, L, P, Q). This phenotype appears to require complete loss of ERK1/2 signaling since overall PV and SST cell densities were unchanged between P60 Cre-negative *Erk1^-/-^* mice and *Erk1^-/-^; Erk2^fl/wt^; Nkx2.1^Cre^* (HET control) animals (**Figure 4-figure supplement 1A**). Additionally, *Erk2^fl/fl^; Nkx2.1^Cre^; Ai14* mutants showed no relative difference in CIN density or subtypes at P30 compared to *Erk2* wildtype controls (**Figure 4-figure supplement 1B-F**). Attempts to generate an *Erk1^-/-^; Erk2^fl/fl^; SST^Cre^* strain yielded no viable mutants at birth (0 of 49, 12 expected) (Taniguchi et al., 2011). Hyperactivation of MEK-ERK1/2 signaling in P21 *Mek1^S217/221E^; Nkx2.1^Cre^; Ai9* mice did not alter the density of SST-expressing CINs, though we observed the previously reported decrease in cortical GABAergic neuron number due to embryonic loss of PV-CINs (**Figure 4-figure supplement 1G-L**) (Holter et al., 2021). Together, these data demonstrate subtype-selective requirements for ERK1/2 signaling in regulating SST expression.

**Figure 4.**
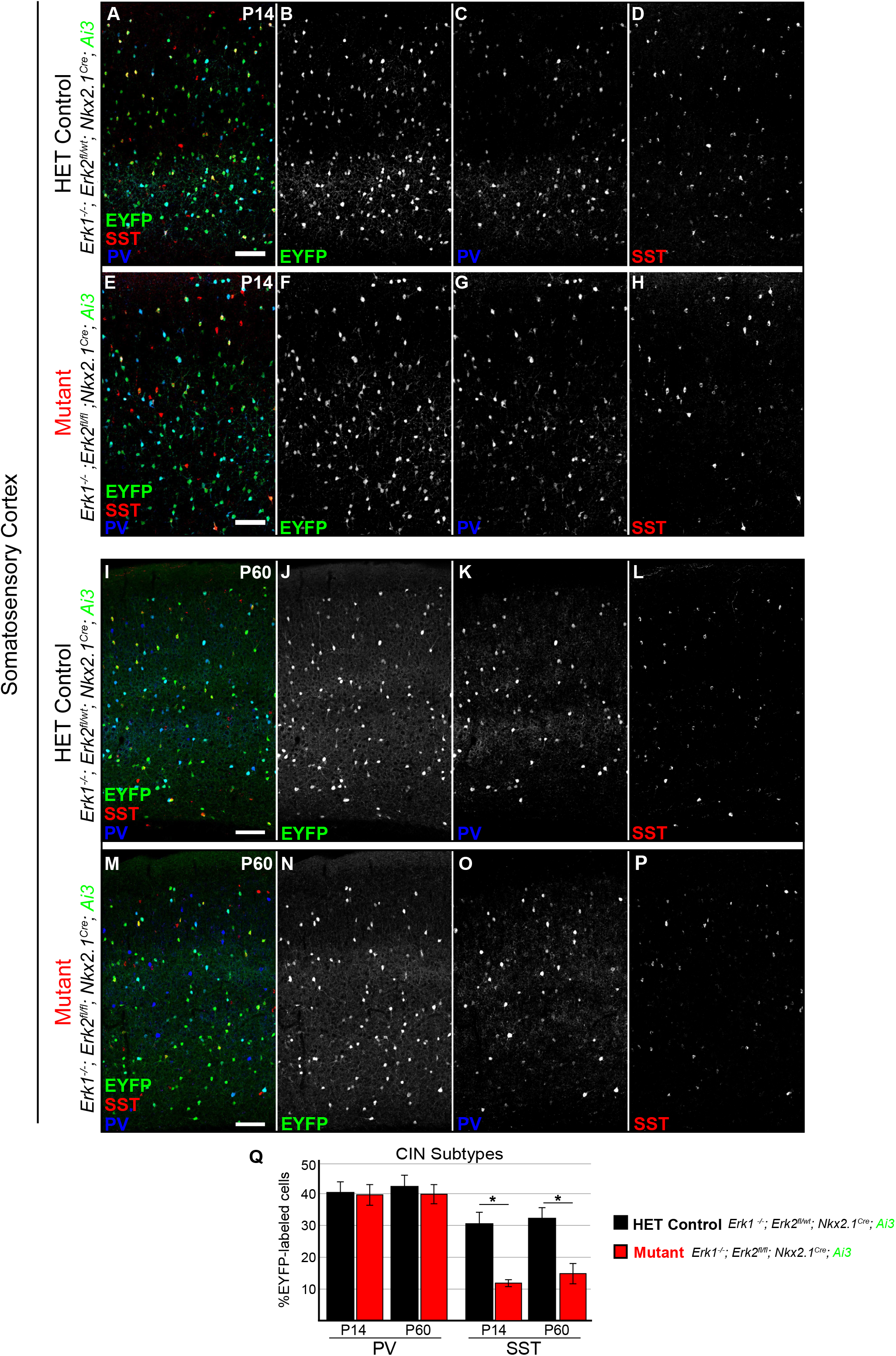
The proportion of SST-expressing CINs decreased following reductions in ERK1/2 signaling. **(A-H)** Representative confocal images of the somatosensory cortex at P14. The proportion of PV-expressing CINs was similar between genotypes (**C, G**, quantification in **Q**; N=3, mean ± SEM, student’s t-test, *p* > .05), while the proportion of SST^+^/EYFP^+^ cells was significantly lower in mutants compared to HET controls (**D, H**, quantification in **Q**; N=3, mean ± SEM, *student’s t-test, *p* = .005). **(I-P)** By P60, no difference in PV-CINs was detected (**K, O**, quantification in **Q**; N=3, mean ± SEM, *p* > .05), whereas a significant decrease in the density of SST^+^/EYFP^+^ cells in mutant cortices was still apparent (**L, P**, quantification in **Q**; N=3, mean ± SEM, *student’s t-test *p* = .03) (scale bars =50 µm).

### Chemogenetic activation of MGE-derived CINs in acute slices

The postnatal maturation of CINs is dependent on trophic cues and neurotransmitter signaling. NRG1/ERBB4, BDNF/TRKB, and glutamate/NMDAR signaling promote appropriate activity-dependent responses in CINs through the activation of multiple intracellular signaling cascades (Fazzari et al., 2010; Huang et al., 1999; Ting et al., 2011; Yap & Greenberg, 2018). However, the specific roles of ERK1/2 signaling in this process remain unclear. To interrogate the cell-autonomous contributions of ERK1/2 to CINs’ activity-dependent gene expression we directly activated a subpopulation of GABAergic neurons using a chemogenetic approach with Designer Receptors Exclusively Activated by Designer Drugs (DREADDs). DREADDs are based on a modified G-Protein-Coupled Receptor and the engineered ligand, clozapine-*N*-oxide (CNO) (Jendryka et al., 2019; Nawaratne et al., 2008; Roth, 2016). In hM3D_q_-DREADD expressing neurons, CNO has been shown to activate G_q_-PLC signaling, resulting in KCNQ channel closure, TRPC channel opening, and ultimately membrane depolarization (Choveau et al., 2012; Mori et al., 2015; Rogers et al., 2021; Roth, 2016; Topolnik et al., 2006; Zou et al., 2016).

We generated *Erk1/2; Nkx2.1^Cre^; G_q_-DREADD-HA* animals for chemogenetic activation of CINs (Rogan & Roth, 2011; Zhu et al., 2016). We performed patch-clamp recordings of FS-CINs in acute slices from P15-P22 HET control and mutant cortices (**Figure 5A**) (Anderson et al., 2010; Kawaguchi, 1997; McCormick et al., 1985). We did not detect a difference in resting membrane potential (CTL -69.00 ± 1.8 mV, MUT -69.75 ± 1.2 mV, *p*=0.73), membrane resistance (CTL 140.0 ± 18 MOhm, MUT 132.5 ± 12 Mohm, *p*= 0.74) or capacitance (CTL 42.5 ± 3.1 pF, MUT 43.5 ± 3.9 pF, *p*=0.85) between genotypes (**Figure 5-figure supplement 1A**). Bath application of 10µM CNO induced a depolarizing increase in the holding current of both HET control and ERK1/2 inactivated FS-CINs (*p*<.05, paired t-test) (**Figure 5B**). Current clamp recordings revealed that depolarizing current injections following application of CNO significantly increased firing frequency when compared to baseline conditions (*p*<.05, paired t-test) (**Figure 5C**). In sum, acute CNO treatment effectively increases the activity of CINs in slices.

**Figure 5.**
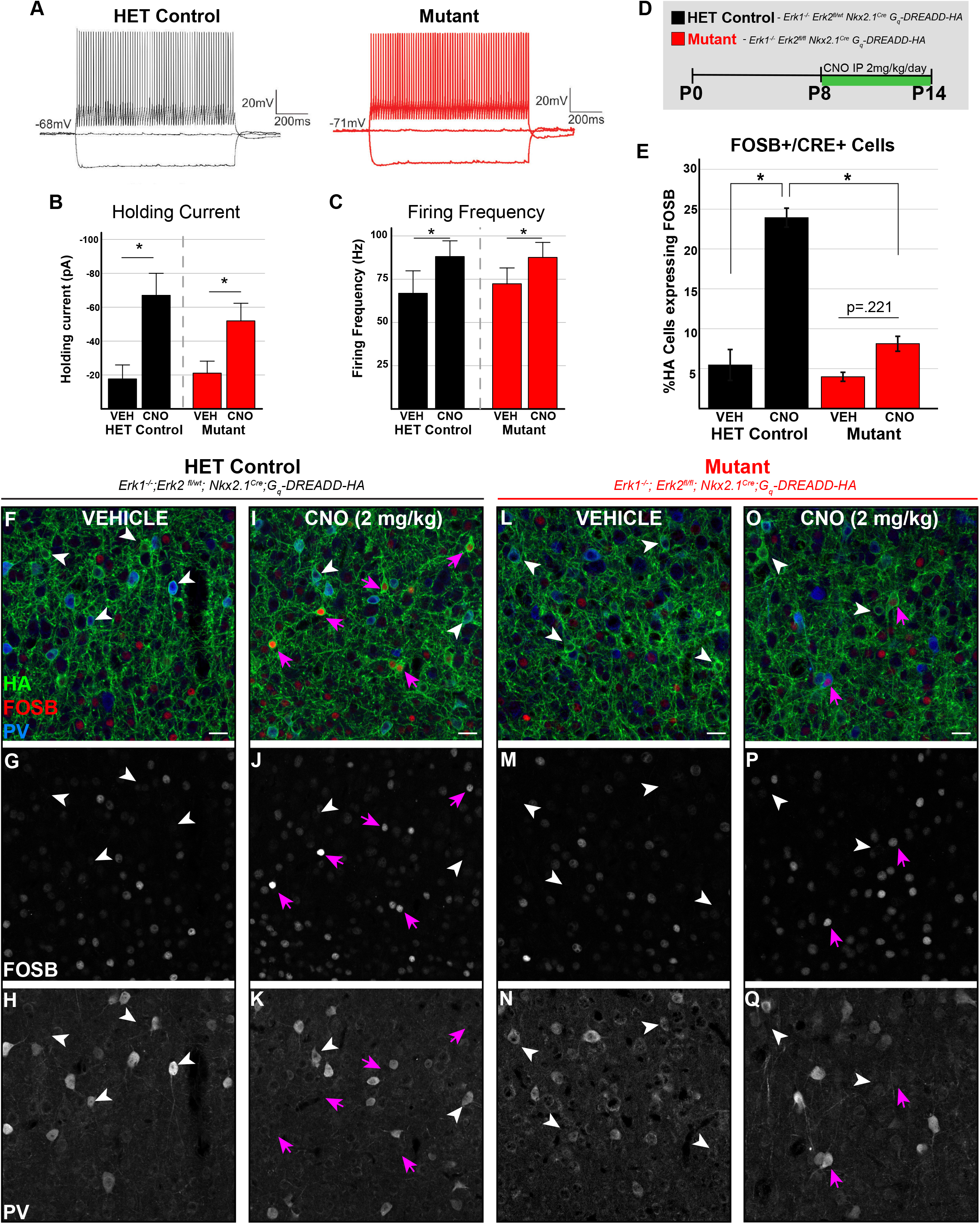
ERK1/2 is required to induce FOSB expression in CINs after h3MD_q_-DREADD mediated stimulation. **(A)** Representative traces of firing response to depolarizing current steps from HET control or mutant neurons indicate that recorded neurons had a fast-spiking electrophysiological phenotype. **(B)** Acute treatment of CINs in acute slices with 10µM CNO led to a depolarizing increase in the holding current in both HET control and mutant neurons (n=8 neurons/genotype, mean +/- SEM, paired t-test, *p* <.05). **(C)** HET control and mutant neurons were recorded under current clamp and the response to a depolarizing current step (200pA, 1s) was measured. CNO significantly increased the firing frequency induced by the applied current step in both HET control and mutant neurons when compared to baseline (VEH) (n= 8 neurons/genotype, mean +/- SEM, paired t-test, *p*<.05). **(D)** Diagram of the in vivo experimental design. Animals were treated with 2 mg/kg CNO or saline vehicle by IP injection for 7 days beginning at P8. **(E)** One week of chemogenetic stimulation increased FOSB expression in HET control CINs but not in mutants (mean +/- SEM, N=4-5/group, one-way ANOVA followed by Fisher’s LSD, *F*_3,14_=15.37, *post-hoc *p*<.001). **(F-Q)** Representative confocal images of FOSB/PV/HA co-expression in layer 5 somatosensory cortex at P14. White arrowheads indicate HA^+^ cells lacking significant FOSB expression, whereas magenta arrows indicate FOSB^+^/HA^+^ co-labeled cells (scale bar = 25 µm).

### ERK1/2 regulates activity-dependent FOSB expression in CINs following chemogenetic depolarization in vivo

During the emergence of GABA_A_R-mediated inhibition in the neonatal cortex, activity is especially critical for the survival of CINs (De Marco García et al., 2011; Hanson et al., 2019; Lodato et al., 2011; Okaty et al., 2009; Pan et al., 2018; Southwell et al., 2012). To assess the role of chronic changes in activity on CIN number in vivo, we treated *Erk1/2; Nkx2.1^Cre^; G_q_-DREADD-HA* animals with saline vehicle or 2 mg/kg CNO daily from P8-P14 (**Figure 5D**). We identified *Nkx2.1^Cre^*-derived CINs using the haemagglutinin (HA) tag fused to the G_q_ receptor in the Cre-dependent construct. The density of Cre-recombined, HA^+^ CINs in the primary somatosensory cortex was not altered by seven days of CNO treatment at this age (**Figure 5-figure supplement 1B**).

In excitatory neurons, ERK1/2 signaling is an important regulator of activity-dependent gene expression (Thomas & Huganir, 2004). However, this relationship is less clear in developing CINs, which exhibit notable differences in activity-dependent gene programs (Hrvatin et al., 2018; Mardinly et al., 2016; Sik et al., 1998). We therefore measured the expression of the activity-dependent, AP-1 family transcription factor, FOSB. Consistent with past studies, we found that 5.4 +/- 1.9% of HA^+^ CINs in vehicle treated, *Erk1^-/-^; Erk2^fl/wt^; Nkx2.1^Cre^; G_q_-DREADD-HA,* HET control mice exhibited detectable FOSB immunoreactivity, whereas we observed many surrounding excitatory neurons with significant FOSB expression (**Figure 5E-H)** (Hrvatin et al., 2018; Mardinly et al., 2016). Vehicle-treated *Erk1^-/-^; Erk2^fl/fl^; Nkx2.1^Cre^; G_q_-DREADD-HA* mutant CINs did not exhibit significant differences in the proportion of FOSB^+^/HA^+^ cells when compared to vehicle treated HET controls (**Figure 5E, L-N**). Lastly, we found that hyperactivation of MEK-ERK1/2 signaling in *Mek1^S217/221E^; Nkx2.1^Cre^; Ai9* mice also did not alter basal FOSB expression in CINs (**Figure 5-figure supplement 1D-F**).

We next analyzed FOSB expression in CINs following one week of chemogenetic stimulation from P8-P14. In HET control mice, CNO led to a significant, 6.3-fold increase in FOSB^+^/HA^+^ co-labeled cells to 23.9 +/- 1.2% of HA^+^ CINs (mean +/- SEM, *F*_3,14_=15.37, post-hoc *p*<.001) (**Figure 5E, I-K, magenta arrows**). At P14, PV^+^/FOSB^+^/HA^+^ triple-labeled cells represented 3.8 +/- 0.4% (mean +/- SEM, N=4) of HA^+^ CINs (**Figure 5I-K**). Importantly, in *Erk1^-/-^; Erk2^fl/fl^; Nkx2.1^Cre^; G_q_-DREADD-HA* mutant CINs, CNO-induced FOSB expression was dramatically reduced when compared to CNO treated HET controls (*F*_3,14_=15.37, post-hoc *p*<.001) (**Figure 5E, O-Q, magenta arrows**). Notably, we were unable to detect a statistically significant increase in FOSB-expressing CINs in CNO treated mutants relative to vehicle treated mutants (post-hoc *p*=.221) (**Figure 5E**). Together, these data demonstrate that ERK1/2 signaling is necessary for chemogenetic activity-dependent FOSB expression in developing CINs.

### Chemogenetic stimulation during development increases SST-expressing CINs in Erk1/2 mutants

Our results suggest ERK1/2 mediates select aspects of activity dependent signaling in CINs. However, chronic chemogenetic activation may be capable of activating ERK1/2-independent pathways that compensate for the effects of *Erk1/2* deletion on SST expression during development. We therefore assessed whether one week of daily CNO treatment between P8 and P14 modified the proportion of SST^+^/HA^+^ cells in the somatosensory cortex. Both calbindin-positive and calbindin-negative SST^+^/HA^+^ populations were decreased in mutant cortices at P14, suggesting a generalized effect of *Erk1/2* deletion on SST-expressing CIN subtypes (**Figure 6A**, SST^+^/CALB1^-^ *p*=.007; SST^+^/CALB1^+^ *p*=.014). In HET control *Erk1^-/-^; Erk2^fl/wt^; Nkx2.1^Cre^; G_q_-DREADD-HA* animals, we found that increasing activity during the second postnatal week did not significantly alter the proportion of SST^+^/HA^+^ CINs (*F*_3,14_ = 22.83, post-hoc *p*=0.144) (**Figure 6B-F**). However, chemogenetic stimulation of *Erk1^-/-^; Erk2^fl/fl^; Nkx2.1^Cre^; G_q_-DREADD-HA* mutants induced a significant increase in SST^+^/HA^+^-CINs relative to vehicle-treated mutants (**Figure 6B, G-J**) (*F*_3,14_ = 22.83, post-hoc *p*=0.005). We did not observe statistically significant changes in the proportion of PV-expressing neurons or co-expression of PV and SST in HA^+^ neurons, indicating that chemogenetic activation did not induce ectopic SST expression in FS CINs (data not shown). The intensity of SST immunoreactivity in CINs reaching a minimum threshold of SST-expression was not significantly different across conditions (**Figure 5-figure supplement 1C**). These data indicate one week of chemogenetic stimulation during early development is sufficient to increase the proportion of CINs expressing SST following the loss of *Erk1/2*.

**Figure 6.**
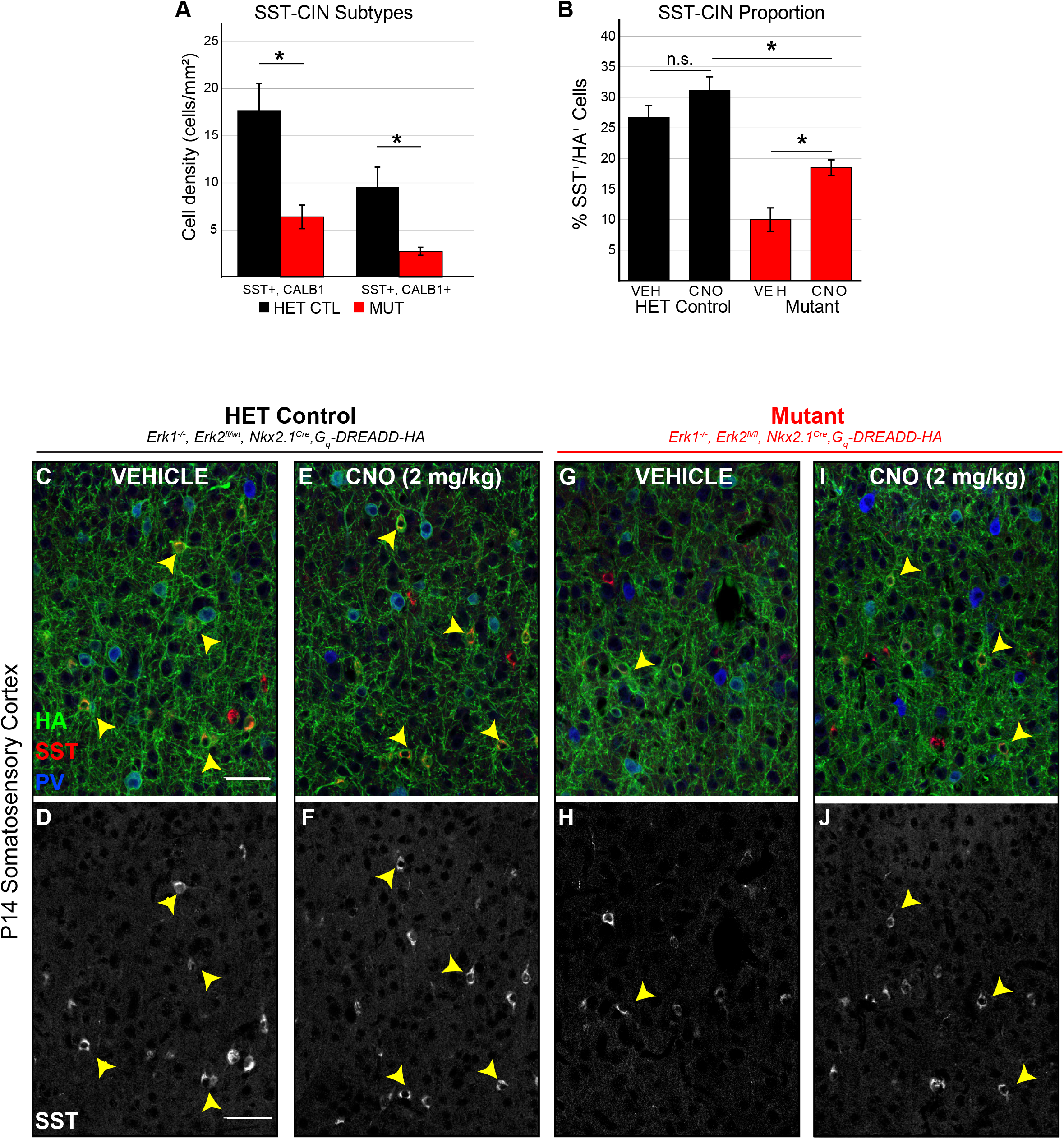
Juvenile chemogenetic stimulation increases SST expression in a subset of CINs. **(A)** Both CALB1+ and CALB1-SST-expressing cells were reduced in P14 mutant animals when compared to HET controls (mean +/- SEM, N=5, student’s t-test, SST+/CALB- *p*=.007, SST+/CALB+ *p*=.014). (**B**) CNO treatment increased the proportion of CINs expressing SST in ERK1/2 inactivated mutants, though this proportion remained significantly lower than in HET control animals (N=4-5, one-way ANOVA followed by Fisher’s LSD, *F*_3,14_=22.83, *post-hoc *p*=.005). **(C-J)** Representative confocal images of CIN subtypes in P14 somatosensory cortex, layer 5 (scale bar = 50µm) (D, F, H, J). Monochrome image highlighting somatostatin immunoreactivity in those cells.

### ERK1/2 signaling is required for activity-dependent FOSB expression in adult CINs

Early cortical circuits may be uniquely capable of responding to altered activity relative to adult stages. We tested the role of ERK1/2 in activity-dependent FOSB expression in adult CINs between 6-10 months of age following one week of daily CNO administration (**Figure 7A**). In HET control adult mice, CNO led to a significant 6.5-fold increase in the proportion of CINs expressing FOSB to 27.6 +/- 2.1% of HA^+^ CINs (mean +/- SEM, *F*_3,8_ = 31.88, post-hoc *p*<0.001) (**Figure 7B-C, J**). At this stage, 16.4 +/- 1.7% of FOSB^+^/HA^+^ CINs expressed PV (mean +/- SEM, N=3). As in young mice, adult CNO-treated *Erk1^-/-^; Erk2^fl/fl^; Nkx2.1^Cre^; G_q_-DREADD-HA* mutant CINs displayed significantly less FOSB induction than HET controls (*F*_3,8_ = 31.88, post-hoc *p*<0.001) (**Figure 7B-E, J**). We noted a reduced proportion of FOSB-expressing HA^+^ CINs in both PV-immunolabeled (4.1 +/- 0.5%) as well as PV-negative subtypes (3.1 +/- 0.7%) (mean +/- SEM, N=3). The overall pattern and number of non-recombined FOSB^+^ cells across the primary somatosensory cortex were relatively unaffected by CIN-specific *Erk1/2* deletion or chronic chemogenetic stimulation (**Figure 7F-I, K**). These data demonstrate that ERK1/2 is required for chemogenetic activation of multiple subtypes of mature CINs.

**Figure 7.**
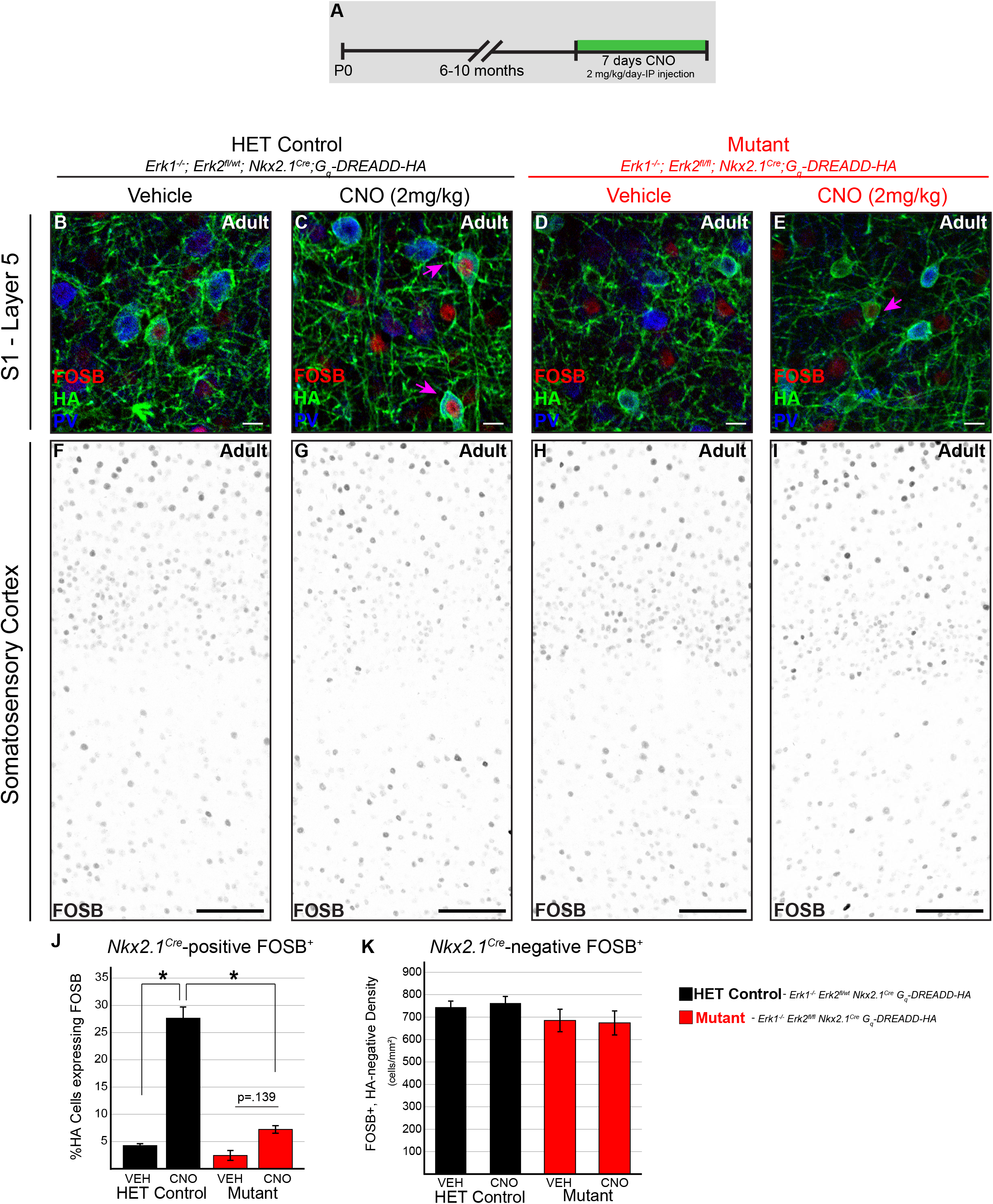
ERK1/2 is required for activity-dependent responses in adults. **(A)** Animals aged 6-10 months were administered 2 mg/kg CNO or saline vehicle via daily IP injection for one week. **(B-E)** High magnification images from layer 5 of the adult somatosensory cortex showing FOSB expression within CINs following 7 days of daily CNO treatment (scale bar = 10µm). One week of CNO treatment increased FOSB expression in HET control CINs but not *Erk1^-/-^; Erk2^fl/fl^; Nkx2.1^Cre^; G_q_-DREADD-HA* mutant CINs (quantification in J). **(F-I)** Confocal images of the adult somatosensory cortex layers 2-5, showing the distribution of FOSB+ cells (scale bar = 100µm). The density of FOSB^+^/*Nkx2.1^Cre^*-negative cells was unchanged in response to *Erk1/2* deletion or chemogenetic stimulation. (**J)** Quantification of *Nkx2.1^Cre^*-positive cells expressing FOSB (N=3, one-way ANOVA followed by Fisher’s LSD, *F*_3,8_ =31.87, *post-hoc *p*<.001). **(K)** Quantification of *Nkx2.1^Cre^*-negative cells expressing FOSB (N=3, one-way ANOVA, *F*_3,8_ = 0.16, *p*>.05).

Finally, we tested whether seven days of chemogenetic stimulation during adulthood altered the proportion of SST-expressing CINs. Following one week of daily CNO treatment, HET control animals showed no change in the proportion of SST^+^/HA^+^ cells in the primary somatosensory cortex (**Figure 8A-F**). In mutant animals, however, we found a modest but statistically significant increase in the proportion of SST^+^/HA^+^ cells following CNO treatment relative to vehicle treated mutants (**Figure 8B, G-J**) (*F*_3,12_ =126.54, post-hoc *p*=0.024). These data suggest that activity-mediated increases in the number of SST-expressing CINs in *Erk1/2* mutants can occur well into adulthood.

**Figure 8.**
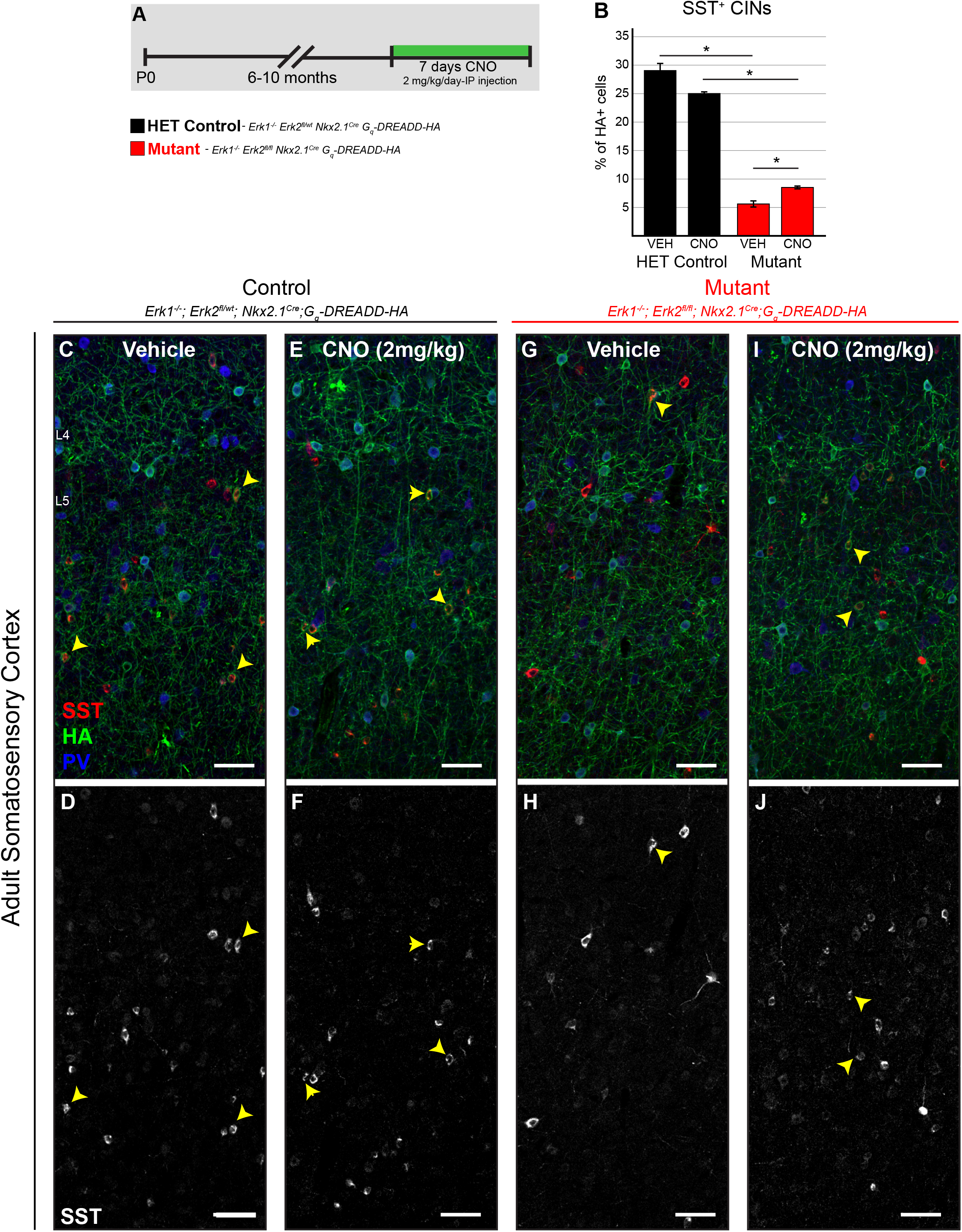
One week of chemogenetic stimulation in adulthood induces a partial rescue of SST expression. **(A)** Diagram of experimental design. **(B)** In adult mice, the proportion of CINs expressing SST is lower in vehicle treated mutants when compared to vehicle treated HET controls. One week of CNO treatment in adult mutants increased SST-CINs relative to vehicle treated mutants (N=4, one-way ANOVA followed by Fisher’s LSD, *F*_3,12_ =126.54, *post-hoc *p*=.024 MUTVEH/MUTCNO). **(C-J)** Confocal images showing immunolabeling for HA, PV, and SST in HET control and *Erk1^-/-^; Erk2^fl/fl^; Nkx2.1^Cre^; G_q_-DREADD-HA* mutant mice. Yellow arrowheads indicate SST^+^/HA^+^ co-labeled cells in layers 4-5 of the primary somatosensory cortex (scale bar =25µm).

## Discussion

Over the course of development, CINs respond to well-defined extracellular cues that regulate proliferation, specification, and the release of neurotransmitters critical for cortical function. Yet the role of associated downstream intracellular signaling cascades in vivo has received less attention. Here, we show that ERK1/2 activity in MGE derivatives is critical for the establishment of glial number but is not necessary for CIN survival or initial commitment to a GABAergic fate. However, we found that ERK1/2 is important for activity dependent expression of FOSB in CINs and regulates the number of SST-expressing, but not PV-expressing, GABAergic neurons. Interestingly, in vivo chemogenetic stimulation of mutant CINs was sufficient to partially overcome the effect of *Erk1/2* inactivation and increase the proportion of CINs expressing SST. These data provide new insight into the specific signaling requirements of developing MGE-derivatives and the cellular functions of ERK1/2 in the development of cortical inhibitory circuits.

Our findings show that loss of ERK1/2 significantly decreases MGE-derived glial number in the anterior commissure, while hyperactive MEK-ERK1/2 signaling is sufficient to increase MGE-derived oligodendrocyte number. Notably, overall oligodendrocyte numbers remained unchanged in ERK1/2 deletion models (**Figure 1–figure supplement 2A**), possibly due to compensatory changes in glia from other progenitor domains (Kessaris et al., 2006; Orduz et al., 2019). These results are consistent with previous findings in neural progenitor domains in the dorsal cortex and peripheral nervous system and further indicate a critical requirement for ERK1/2 signaling in establishing myelinating glial number across the nervous system, despite substantial differences in origin and local extracellular environments (Dinh Duong et al., 2019; Filges et al., 2014; X. Li et al., 2012; Newbern et al., 2011). Whether the reduction in MGE-derived glial number is due to a disruption in the gliogenic transition in neural progenitors or subsequent stages of glial development is unknown (Ishii et al., 2012; S. Li et al., 2014; X. Li et al., 2012), but our results are consistent with a key role for FGF-ERK1/2 signaling in driving forebrain oligodendrocyte development (Furusho et al., 2011, 2017; Ishii et al., 2016). It will be interesting to further dissect the specific molecular mechanisms of ERK1/2 contributions to glial development with Cre lines that recombine at different stages.

Upstream RASopathy mutations in *Ras* and *Nf1* alter multiple parallel kinase cascades, including ERK1/2, increase glial number, and alter white matter structure (Angara et al., 2020; Breunig et al., 2015; Fattah et al., 2021; Gutmann et al., 2001; Holter et al., 2019; Krencik et al., 2015; López-Juárez et al., 2017; Rizvi et al., 1999). Whether reduced *ERK1/MAPK3*, *ERK2/MAPK1,* or *MEK2/MAP2K2* gene dosage in individuals with 16p11.2-, distal22q11.2-, or 19p13.3-microdeletions, respectively, is sufficient to disrupt myelinating glial number and contribute to abnormal white matter microstructure observed in some of these conditions is unclear (Chang et al., 2016; Owen et al., 2014; Qureshi et al., 2014; Silva et al., 2022). This conserved functional link may also be an important consideration in pediatric RASopathy clinical trials utilizing prolonged pharmacological ERK1/2 pathway inhibition (Payne et al., 2019).

Past studies have shown that distinct neuronal populations arising from the same progenitor domain exhibit unexpectedly diverse cellular responses to ERK1/2 modulation (Holter et al., 2021; Hutton et al., 2017; Newbern et al., 2011; Sanchez-Ortiz et al., 2014; Vithayathil et al., 2015; L. Xing et al., 2016). In many non-neuronal cells, ERK1/2 and PI3K/AKT are important regulators of proliferation, migration, and survival (Lavoie et al., 2020). In CINs, PI3K/AKT mediates comparable cellular behaviors (Oishi et al., 2009; Polleux et al., 2002; Vogt et al., 2015; Wei et al., 2020; Wong et al., 2018). Early studies in cancer cell lines suggested a critical role for ERK1/2 in migration (Klemke et al., 1997) and an early CNS-wide *Erk1/2* conditional knockout exhibited reduced calbindin- and GAD67-expressing, presumptive GABAergic neurons in the P0.5 cortex (Imamura et al., 2010). In contrast, pharmacological MEK1/2 inhibitors did not disrupt embryonic CIN migration in acute slice assays (Polleux et al., 2002). We used two strains of mice with improved selectivity for the GABAergic lineage to show that ERK1/2 is not necessary for CIN migration or survival in vivo. This outcome differs from our previously reported requirement for ERK1/2 signaling in excitatory corticospinal neuron survival during the neonatal period (L. Xing et al., 2016). The mouse models we have generated will provide important tools for further dissection of the precise mechanisms that regulate these cell context-dependent functions.

Early CIN development is determined, in part, by local morphogen gradients acting via specific transcriptional programs (Hu et al., 2017; Lim et al., 2018). For example, SHH signaling is required for the specification of MGE-derived GABAergic neurons during mid embryogenesis and later biases these neurons toward a SST-expressing fate (Tucker et al., 2008; Tyson et al., 2015; Xu et al., 2005, 2010; Yu et al., 2009). Moreover, the pharmacological MEK1/2 inhibitor, U0126, blocks the SHH-induced production of OPCs in cultured dorsal neural progenitors (Kessaris et al., 2004). In certain contexts, ERK1/2 signaling has been shown to non-canonically activate GLI1, a well-known transcriptional target of SHH (Pietrobono et al., 2019; Po et al., 2017). It is possible that SHH and ERK1/2 signaling converge on GLI1 to direct neurons toward a SST-expressing CIN fate. Indeed, conditional loss of *Nf1* or expression of a hyperactivating *BRaf^V600E^* variant leads to a reduction in PV-CINs and an increase in the proportion of SST-expressing CINs (Knowles and Stafford et al., 2022). SST expression alone provides an incomplete view of the RS-CIN lineage. Examination of a panel of genes associated with SST-expressing CINs that includes ion channels, transcription factors, and synaptic molecules alongside SST would be informative. Future studies will be necessary to identify the molecular mediators associated with changes in ERK1/2 activity and CIN specification in the developing CNS.

Cortical circuit formation establishes a complex balance between GABAergic and glutamatergic signaling. GABA-mediated signals promote the maturation of excitatory neurons, in part via ERK1/2 dependent mechanisms (Cancedda et al., 2007; Obrietan et al., 2002; Peerboom & Wierenga, 2021). Past reports demonstrate that early alterations in CIN number or activity can permanently change circuit architecture and animal behavior (Bitzenhofer et al., 2021; Kaneko et al., 2022; Magno et al., 2021; Tuncdemir et al., 2016). Here, we employed a chemogenetic approach to selectively activate CINs with temporal specificity. We found that *Nkx2.1^Cre^*-targeted chemogenetic stimulation during the second postnatal week did not alter CIN number. Past studies have identified a role for activity in early CIN survival (Denaxa et al., 2018; Priya et al., 2018; Southwell et al., 2012; Wong et al., 2018). However, we modulated the activity of endogenous CINs at a stage past the reported peak of CIN apoptosis and after GABA_A_Rs acquire inhibitory responses (Ben-Ari et al., 2007, p.; Denaxa et al., 2018; Southwell et al., 2012; Wong et al., 2018). It is not clear whether CIN-specific ERK1/2 signaling is required for global cortical circuit activity or cognitive behaviors (Adler et al., 2019; Chen et al., 2019; Cummings & Clem, 2020; Fu et al., 2014; Kluge et al., 2008; C. Lee et al., 2022; Magno et al., 2021).

Recent work has led to a deeper appreciation for the unique contributions of CIN subtypes to cortical microcircuits. While our past studies have clearly identified significant effects of MEK-ERK1/2 hyperactivation on PV-CIN lineage specification, we failed to observe changes in PV-CIN number, intrinsic membrane properties, or the expression of PV following *Erk1/2* deletion (Angara et al., 2020; Holter et al., 2021; Knowles et al., 2022). However, we now show that ERK1/2 signaling is necessary for activity-dependent FOSB expression in adult PV-CINs.

Unexpectedly, we found that ERK1/2 is critical for SST protein expression in a subset of CINs. Somatostatin is developmentally regulated; *Sst* mRNA is first detected in immature CINs by mid-embryogenesis, but substantial levels of SST protein are not observed in CINs until early neonatal stages and patterns of SST expression do not fully mature until P30 (Batista-Brito et al., 2008; Denaxa et al., 2012; Forloni et al., 1990; D. R. Lee et al., 2022; Ma et al., 2022; Neves et al., 2013; Taniguchi et al., 2011). A loss of SST protein has been seen in a number of other neuropathological conditions, but has not been directly assessed in conditions characterized by reduced ERK1/2 signaling (Davies et al., 1980; Fee et al., 2017; Pantazopoulos et al., 2017; Peng et al., 2013; Robbins et al., 1991; Sun et al., 2020; Tripp et al., 2011; Wengert et al., 2021). SST is not only a marker of RS-CINs, but an active neuropeptide with biological functions (Baraban & Tallent, 2004; Grilli et al., 2004; Liguz-Lecznar et al., 2016; Smith et al., 2019; Song et al., 2021; Tereshko et al., 2021). The release of bioactive neuropeptides from subtypes of CINs likely contributes to cortical computation (Ouwenga et al., 2018; Smith et al., 2019). SST has an overall inhibitory role on cortical circuits and loss of SST protein can cause subtle deficits in motor learning (Tallent & Qiu, 2008; Zeyda et al., 2001). Indeed, a recent report demonstrated that exogenous SST infusion into V1 can improve performance in a visual discrimination task, independent of GABA (Song et al., 2020). Administration of SST or synthetic analogues may compensate for the aberrant development of SST-expressing CINs in *Erk1/2* loss of function models (Banks et al., 1990; Engin & Treit, 2009; Kiviniemi et al., 2015; McKeage, 2015). Approaches that target SST may have broader applications in the treatment of neurodevelopmental disorders, particularly select cases of epilepsy, autism, and schizophrenia where deficits in SST-CIN function are well described (Song et al., 2021).

Interestingly, one week of chemogenetic stimulation partially restored SST expression in *Erk1/2*-deleted CINs at both juvenile and adult stages. Chemogenetic depolarization may engage several parallel cascades including cAMP/PKA, Ca^2+^, PKC, and ERK1/2, which often converge on common transcriptional targets, such as CREB, SRF, and AP-1 family members (Yap & Greenberg, 2018). We demonstrated that the activity-dependent expression of FOSB, a CREB target, requires ERK1/2 signaling in CINs, similar to glutamatergic neurons (Guenthner et al., 2013; Hrvatin et al., 2018; Mardinly et al., 2016; Spiegel et al., 2014). Reduced SST expression in a subset of CINs in *Erk1/2* deleted animals limited our ability to directly assess whether ERK1/2 signaling is required for FOSB expression in this specific CIN subtype. Interestingly, one of the first CREB target genes to be identified was SST (Montminy & Bilezikjian, 1987; Yamamoto et al., 1988). Chemogenetic activation of ERK1/2-independent cascades appears sufficient to partially compensate for the loss of *Erk1/2* and modestly increase SST expression in CINs. Whether other activity-dependent genes critical for CIN plasticity are similarly modified remains to be seen. Our finding that adult chemogenetic activation partially restored SST expression raises the possibility that adult modulation of neural activity may be capable of reversing select developmental deficits in ERK1/2-related syndromes.

## Acknowledgements

We thank Chris Wedwick, Anna Bayne, Mya Breitweiser, and Elise Bouchal for technical assistance with mouse sample processing and image analysis. This research is supported by the National Institute of Health grants R00NS076661 and R01NS097537 awarded to JMN. SK is supported by the ARCS Foundation. AMS and DV are supported by the Spectrum Health-Michigan State Alliance Corporation, Autism Research Institute pilot grant and Department of Defense grant NF200109. TA and GL are supported by the University of Arizona – COM Phoenix “Springboard” award. We deeply appreciate the support of the ASU Keck Bioimaging facility and ASU Personalized Diagnostic Core for outstanding technical advice and support.

## Materials and Methods

### Genetically modified mice

All mice were fed *ad libitum,* housed under a 12-hour light-dark cycle, and studied in accordance with ARRIVE and Institutional Animal Care and Use Committee guidelines at Arizona State Univ., Michigan State Univ., and the Univ. of Arizona. Mature mice examined in this study were euthanized via CO2 inhalation or approved chemical anesthetics prior to transcardial perfusion, while embryonic/neonatal mice were cryoanesthetized, as described in the AVMA Guidelines on Euthanasia. Male and female mice on mixed genetic backgrounds were utilized in all experiments in this work and generated from crosses between the following individual strains of mice.

| Mouse Strains | Details | Reference | Identifier |
| --- | --- | --- | --- |
| *Erk1/Mapk3*<sup>-/-</sup> | Neo cassette inserted into Exon 1-6 of *Erk1/Mapk3* | Selcher et al, 2001 | RRID:IMSR_JAX:019113 |
| *Erk2/Mapk1*<sup>loxp/loxp</sup> | *Erk2/Mapk1* exon 2 flanked by loxp (“*fl*”) | Samuels et al., 2008 | RRID:IMSR_JAX:019112 |
| *CAG-LSL-Mek1*<sup>S217/221E</sup> | Cre-dependent hyperactive *Mek1*<sup>S217/221E</sup> overexpressing mice | Krenz et al., 2008 | MGI:3822131 |
| *Nkx2.1*<sup>Cre</sup> | Cre recombinase expression in radial glia in the MGE, POA, and other regions by E8.5 | Xu et al., 2008 | RRID:IMSR_JAX:008661 |
| *Slc32a1/VGAT*<sup>Cre</sup> | Cre recombinase expression in all postmitotic GABAergic neurons, including CINs by E11.5 | Vong et al., 2011 | RRID:IMSR_JAX:016962 |
| *CAG-LSL-Gq-DREADD* | Cre-dependent HA-hM3Dq-pta-mCitrine expressing line | Zhu et al., 2016 | RRID:IMSR_JAX:026220 |
| *Rpl22*<sup>HA</sup> | Cre-dependent RNA immunoprecipitation line | Sanz et al., 2009 | RRID:IMSR_JAX:011029 |
| *Ai3 / Ai9 / Ai14* | Cre-dependent fluorescent protein (EYFP or tdTomato) expressing lines | Madisen et al., 2010 | RRID:IMSR_JAX:007903, 007908, 007903 |

To generate *Erk1/Erk2* loss-of-function mutants, mice expressing *Nkx2.1^Cre^* or *Slc32A1/VGAT^Cre^* were bred with mice possessing a neo-insertion in exons 1-6 in the *Erk1/Mapk3* gene (referred to as *Erk1^-^*) and/or a loxp flanked exon 2 in the *Erk2/Mapk1* gene (referred to as *Erk2^fl^* ) (Monory et al., 2006; Samuels et al., 2008; Selcher, 2001; Xu et al., 2008). Littermates expressing Cre recombinase and heterozygous *Erk2^fl/wt^* were often used as controls for complete *Erk1/2* loss-of-function experiments unless stated otherwise. Cre-dependent tdTomato/RFP (*Ai9 or Ai14*) or eYFP (*Ai3*) strains were employed to fluorescently-label Cre-expressing cells (Madisen et al., 2010).

Conditionally hyperactive MEK-ERK1/2 mutants were generated by crossing *Nkx2.1:Cre* mice with *CAG-loxp-STOP-loxp-Mek1^S217/221E^*(referred to as *Mek1^S217/221E^)* mice on a mixed genetic background to generate double heterozygous mutants expressing *Mek1^S217/221E^* in a Cre-dependent pattern. *Mek1^S217/221E^*mice were kindly provided by Dr. Maike Krenz and Dr. Jeffrey Robbins (Krenz et al., 2008).

For chemogenetic experiments, an *loxp-STOP-loxp-G_q_-DREADD-HA* line was used, which produces the engineered G-protein coupled receptor, hM3D_q_, with a hemagglutinin (HA) tag in Cre-expressing cells (Zhu et al., 2016). To chemogenetically stimulate these neurons in vivo, mice of the indicated age were randomly assigned to receive an intraperitoneal injection of 2 mg/kg body weight clozapine-*N*-oxide (CNO) or 0.09% NaCl saline vehicle every 24 hours for 7 consecutive days. Body weight and time of injection were recorded each day and tissue was collected 2 hours after the last dose of CNO.

### PCR Genotyping

Genomic DNA was rapidly extracted from mouse tissue samples for PCR genotyping with 25mM NaOH, 0.2 mM EDTA, pH=12 and subsequently neutralized with 40 mM Tris-Cl, pH=5. The primers used for genotyping were as follows: (listed 5’-3’): *Cre* – TTCGCAAGAACCTGATGGAC and CATTGCTGTCACTTGGTCGT to amplify a 266 bp fragment; *Erk1^-^* – AAGGTTAACATCCGGTCCAGCA, AAGCAAGGCTAAGCCGTACC, and CATGCTCCAGACTGCCTTGG to amplify a 571 bp segment wildtype and a 250 bp segment KO allele; *Erk2^fl^* – AGCCAACAATCCCAAACCTG, and GGCTGCAACCATCTCACAAT amplify 275 bp wildtype and 350 bp floxed alleles; *Ai3/Ai9* – four primers were used - AAGGGAGCTGCAGTGGAGTA, CCGAAAATCTGTGGGAAGTC, ACATGGTCCTGCTGGAGTTC, and GGCATTAAAGCAGCGTATCC amplify a 297 bp wildtype Rosa26 segment and a 212 bp *Ai3*/*Ai9* allele; *hM3Dq* – CGCCACCATGTACCCATAC and GTGGTACCGTCTGGAGAGGA amplify a 204 bp fragment; *Mek1^S217/221E^* – GTACCAGCTCGGCGGAGACCAA and TTGATCACAGCAATGCTAACTTTC amplify a 600 bp fragment.

### Tissue Preparation and Immunostaining

Mice of the specified age were fully anesthetized and perfused with cold 1X PBS followed by 4% PFA in a 1X PBS solution. Brains were then removed and post-fixed at 4C. Brains were sliced with a vibratome or were cryoprotected for preparation on a cryostat. Sectioned brain tissue was then incubated for 24-48 hours at 4°C in a primary antibody solution diluted in .05 – 0.2% Triton in 1X PBS with 5% Normal Donkey Serum (NDS). Primary antibody information is provided in the table below.

Sections were then washed three times in 1X PBS .05% Triton and incubated in a solution containing secondary antibodies diluted to 1:1000 in 1X PBS .05 – 0.2% Triton and 5% NDS. A Zeiss LSM710 or LSM800 laser scanning confocal microscope was used to collect images. Representative confocal images were optimized for brightness and contrast in Adobe Photoshop and compiled into figures with Adobe Illustrator.

| Primary Antibody and Host | Source | Identifier |
| --- | --- | --- |
| Rabbit Anti-Somatostatin | Peninsula/BMA Biomed. | Cat# T-4103.0050, RRID:AB_518614 |
| Guinea Pig Anti-Somatostatin | Synaptic Systems | Cat# 366 004, RRID:AB_2620126 |
| Goat Anti-Parvalbumin | SWANT | Cat# PVG-213, RRID:AB_2650496 |
| Chicken Anti-GFP | Aves | Cat# GFP-1020, RRID:AB_10000240 |
| Chicken Anti-RFP | Rockland | Cat# 600-901-379, RRID:AB_10704808 |
| Rabbit Anti-RFP | Rockland | Cat# 600-401-379, RRID:AB_2209751 |
| Rabbit Anti-ERK2 | Abcam | Cat# ab32081, RRID:AB_732210 |
| Chicken Anti-Calbindin 1 | Novus | NBP2-50028 |
| Mouse Anti-NeuN | Millipore | Cat# MAB377, RRID:AB_2298772 |
| Mouse Anti-HA | Biolegend | Cat# 901513, RRID:AB_2565335 |
| Rabbit Anti-FOSB | Abcam | Cat# ab184938, RRID:AB_2721123 |
| Rabbit Anti-MEK1 | Abcam | Cat# ab32091, RRID:AB_776273 |
| Rabbit Anti-GFAP | Abcam | Cat# ab7260, RRID:AB_305808 |
| Rabbit Anti-OLIG2 | Millipore | Cat# AB9610, RRID:AB_570666 |

### Image Analysis

For all immunolabeling experiments, images from multiple tissue sections per mouse containing the anatomical region of interest were quantified by an observer blinded to the genotype. At least three biological replicates per genotype were derived from at least three independent litters to minimize litter batch effects and littermate controls were frequently utilized. To estimate labeled cell density, regions of interest were defined using standard anatomical and architectonic landmarks, the area was measured in Photoshop, and labeled cells were quantified. Sampling information for assessments of anterior commissure glia can be found in Figure 1 – figure supplement 2A. Cortical density measurements are derived from counts collected across an entire cortical column. For analyses of postnatal samples, total CIN density and co-labeled proportions were measured from a minimum of three mice per genotype and at least 500 Cre-expressing CINs derived from confocal images of at least three sections per mouse. Power analyses were performed prior to conducting most experiments to estimate sample size using a significance level of 5%, power of 80%, and standard deviation based on pilot studies or previously published work. For quantitative analysis of the intensity of SST immunoreactivity in CINs, we analyzed images which were acquired with the same settings across trials. We sampled from a minimum of 10 randomly selected cells per animal that met a minimal threshold for SST-expression and three animals per condition. Integrated density of SST immunoreactivity was measured using Photoshop. For comparisons involving two groups, statistical significance was determined using an unpaired, two-tailed t-test. For comparison of three or more groups, we performed a one-way ANOVA followed by Fisher’s LSD post-hoc tests using IBM SPSS Statistics 28.

### RNA collection and sequencing

RNA collection and ribosome immunoprecipitation from brain samples was performed essentially as described in Sanz et al., 2009. Whole P7.5 cortices were rapidly dissected, rinsed, and Dounce homogenized in 500 µL of ice-cold RNase-free buffer polysome buffer (50 mM Tris pH 7.5, 100 mM KCl, 12 mM MgCl2, 1% NP-40, 1 mM DTT, 200U/ml Promega RNasin, 1 mg/ml heparin, 100 µg/ml cycloheximide, and protease inhibitors). The lysate was cleared via centrifugation at 10,000g for 20 min at 4°C. 60 µl of supernatant (“INPUT”) containing RNA derived from the entire cortical sample was collected and lysed with 350 µl of buffer RLT from Qiagen RNeasy Kit supplemented with 10μl/ml bME. Rpl22^HA^*-*containing ribosomes were immunoprecipitated (“IP”) from the remaining supernatant by gently mixing with 5µl of rabbit anti-HA antibody (#71–5500 Thermo Fisher Sci) and incubating with gentle rocking at 4°C for 2 hours before isolation with 50 µl of protein A/G magnetic beads previously equilibrated in polysome buffer and a magnetic stand. Immunoprecipitates were rinsed 3x with 800 µl of a high salt buffer (50 mM Tris pH 7.5, 300 mM KCl, 12 mM MgCl2, 1% NP-40, 1 mM DTT, 100 µg/ml cycloheximide) and then lysed in 350 µl of buffer RLT. RNA isolation from INPUT and IP fractions was performed following manufacturer’s instructions using Qiagen’s RNeasy extraction kit with DNAse digestion. RNA quantitation was performed using a Ribogreen RNA reagent and samples with an RNA integrity number < 7 on an Agilent Bioanalyzer were selected for sequencing.

2 ng of purified RNA were used for single primer isothermal amplification (SPIA) and generation of double-stranded cDNA with the Ovation RNAseq System V2 per manufacturer instructions (Nugen Technologies). Library construction and sequencing were performed on the Illumina NextSeq 500 platform at the Arizona State University’s Genomics Core facility. RNAseq reads were quality checked, filtered, aligned to the mouse genome Ensembl GRCm38 primary assembly using RNA-STAR, and read counts were estimated using featureCounts (Dobin et al., 2013). DEXseq was employed to measure differential exon usage in *Erk1* exons 1-6 and *Erk2* exon 2 (Anders et al., 2012). Analysis with DeSeq2 resulted in >11,000 protein-coding genes in each sample with a normalized count > 50 (**Table S1**) (Love et al., 2014). Protein-coding genes were considered differentially expressed if the DeSeq2 derived fold-change between conditions was >1.5, unadjusted p-value <.05, and the CoV of normalized DeSeq2 counts within a condition were < 90 (**Table S2**). All code is publicly available, please see original papers for full information. Raw sequencing data are available in NCBI’s Gene Expression Omnibus repository at accession number GSE206633 (Edgar et al., 2002).

### Electrophysiology

For electrophysiological experiments, postnatal day 15-22 (P15-22) mice were used for the preparation of in vitro whole-cell patch-clamp experiments as previously reported (Anderson et al., 2010; Goddeyne et al., 2015; Holter et al., 2021; Nichols et al., 2018). The somatosensory cortices of these mice were injected with a Cre-dependent AAV9-FLEX-tdTomato (Addgene catalog #28306-AAV9) viral vector solution between P0-P2 to unambiguously label a subset of recombined CINs for fluorescently-guided patch clamping. In brief, mice were then deeply anesthetized by isoflurane inhalation before decapitation. Brains were quickly removed and immersed in the saturated (95% O_2_, 5% CO_2_), ice-cold artificial cerebral spinal fluid (aCSF), which contains (in mM): 126 NaCl, 26 NaHCO_3_, 2.5 KCl, 10 Glucose, 1.25 Na_2_H_2_PO_4_H_2_O, 1 MgSO_4·_7H_2_O, 2 CaCl_2_H_2_O, pH 7.4. Coronal slices from the region of the somatosensory cortex (350 μm in thickness) were collected with a vibratome (VT 1200; Leica, Nussloch, Germany). Following the preparation of brain slices, they were allowed to recover in the same aCSF at 32°C for 30 minutes before being moved to room temperature for an additional 30 minutes. For patch-clamp recordings, slices were transferred into a submerged recording chamber and perfused continuously with aCSF at 32°C at a rate of 1-2 ml/min. Whole-cell patch-clamp recordings were obtained from fluorescent positive neurons in layer V (L5) of the somatosensory cortex by using an Axon 700B amplifier. The data were filtered at 2 kHz and sampled at 10 kHz via a digitizer (Digidata 1440, Molecular Devices) and recorded using Clampex 10.6 software (Molecular Devices). The pipettes had a resistance of 2-5 Mῼ when filled with the internal solution which contains (in mM): 135 K-Gluconate, 4 KCl, 2 NaCl, 10 HEPES, 4 EGTA, 4 Mg ATP and 0.3 Na Tris. Pipettes were pulled from borosilicate glass (BF150-110-10, Sutter Instruments) by using a Sutter Instrument puller (Model P-1000). The stability of the recordings was monitored throughout the experiment and recordings with series resistances (Rs) larger than 25 Mῼ or a change of more than 20% were abandoned.

Data were analyzed using Clampfit software (Molecular Devices) and results were presented as mean ± SEM. Membrane resistance (Rm) and capacitance (Cm) were calculated from the response to a voltage step (5mV). Resting membrane potential was measured during current-clamp recording prior to the introduction of steps (-200pA, 0 or 200pA, 1 s duration) used in examining the firing properties of the neurons. For calculation of CNO induced changes in holding current, neurons were recorded under voltage clamp at -70mV. The change in holding current was calculated by subtracting the baseline value (i.e., in aCSF) from that measured after 20 minutes of bath application of CNO (10 µM). At this time point, CNO-induced changes in holding current had reached a steady-state plateau. Data were compiled and appropriate statistical analysis was performed using Prism software (Graphpad) and figures compiled in CorelDraw (Corel Corporation) and Adobe Illustrator. For all electrophysiological statistics, *p*< .05 was considered statistically significant and tested with an unpaired student t-test unless otherwise indicated.

---

**Figure 1 - figure supplement 1.**
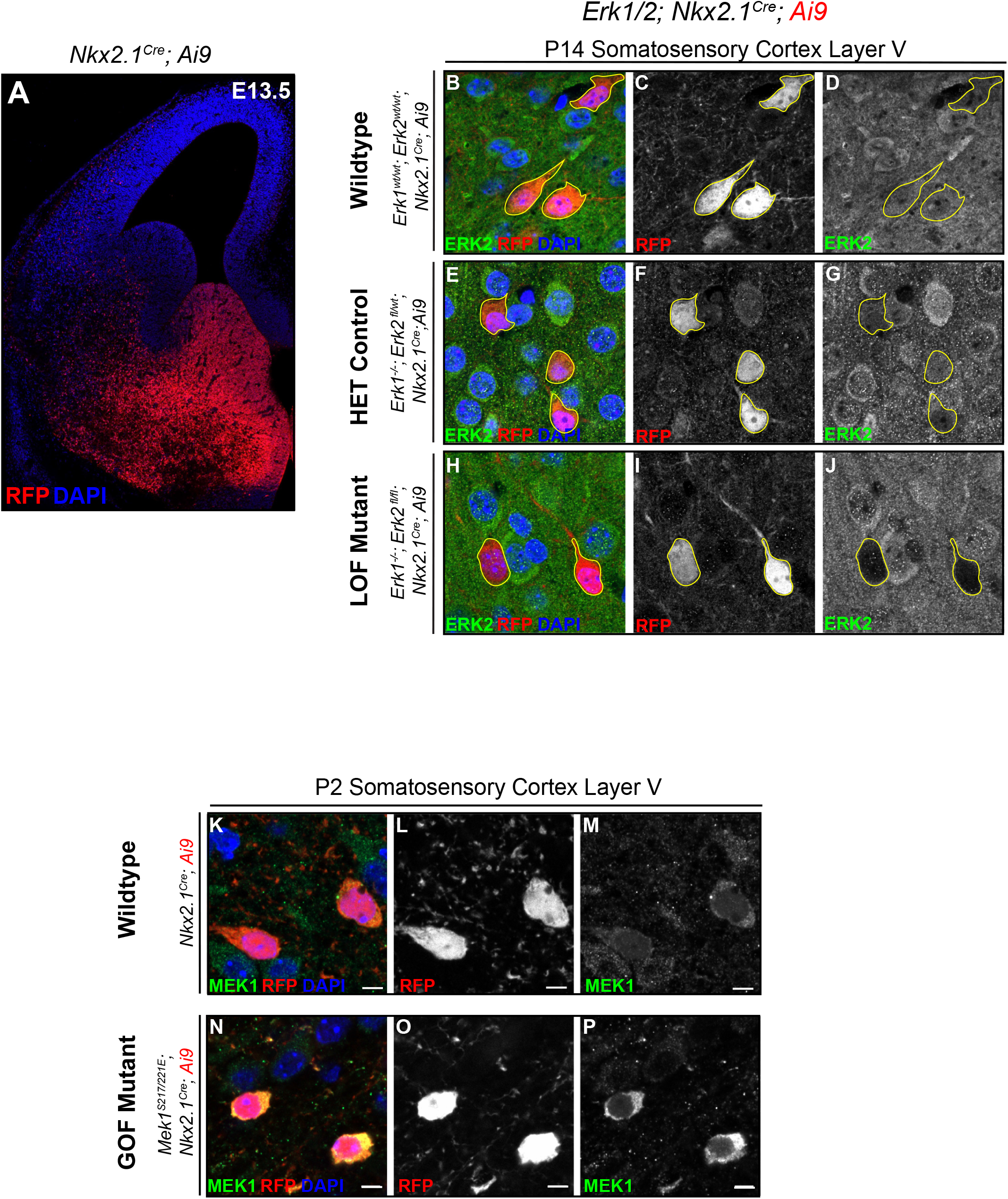
Cre-targeted modulation of ERK1/2 signaling. **(A)** Coronal hemisection of an E13.5 *Nkx2.1^Cre^; Ai9* brain showing the expected pattern of *Nkx2.1^Cre^*-mediated recombination. **(B-J)** High-resolution confocal images of neurons demonstrating reduced ERK2 immunoreactivity in RFP-expressing cells in P14 *Erk1^-/-^; Erk2^fl/fl^; Nkx2.1^Cre^; Ai9* mice (H-J) compared with wildtype (B-D) or heterozygote controls (E-G) (scale bar = 5µm). **(K-P)** High-resolution confocal images demonstrating an increase in MEK1 expression in RFP-expressing neurons in *Mek1^S217/221E^*; *Nkx2.1^Cre^; Ai9* (N-P) animals compared to wildtype (K-M) (scale bar = 5µm).

**Figure 1 - figure supplement 2.**
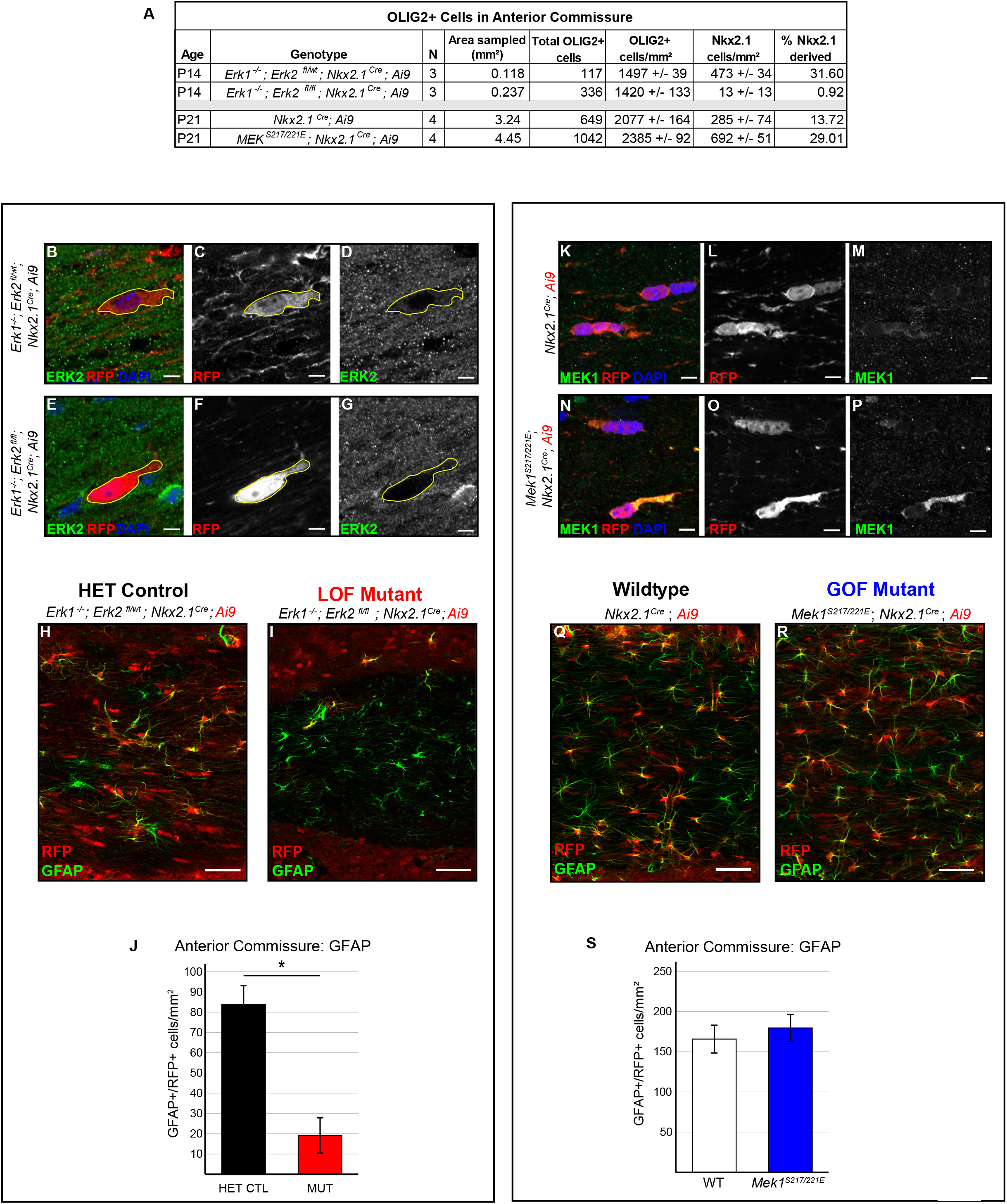
Reduced ERK1/2 signaling regulates MGE-derived GFAP^+^ astrocyte number. **(A)** Extended data on *Nkx2.1^Cre^*-derived OLIG2^+^ cells in the anterior commissure. **(B-G)** High-resolution confocal images demonstrating reduced ERK2 immunoreactivity in RFP-expressing cells in the anterior commissure of P14 *Erk1^-/-^; Erk2^fl/fl^; Nkx2.1^Cre^; Ai9* mice (E-G) compared with heterozygote controls (B-D) (scale bar = 5µm). **(H-J)** Images demonstrating reduced density of RFP^+^/GFAP^+^ co-labeled cells in the anterior commissure of *Erk1^-/-^; Erk2^fl/fl^; Nkx2.1^Cre^; Ai9* mice compared with heterozygous controls. (Quantification in **J**, N=3, mean ± SEM, *student’s t-test, *p*<.05) (scale bar = 50µm). **(K-P)** High-resolution confocal images demonstrating an increase in MEK1 protein expression in RFP-expressing neurons in *Mek1^S217/221E^*; *Nkx2.1^Cre^; Ai9* animals (N-P) compared to wildtype controls (K-M) (scale bar = 5µm). **(Q-S)** Images of RFP+/GFAP+ cells in the anterior commissure of wildtype and *Mek1^S217/221E^*; *Nkx2.1^Cre^; Ai9* mice compared with heterozygous controls. (Quantification in **S**, N=3-4, mean ± SEM, *student’s t-test, *p*>.05) (scale bar = 50µm).

**Figure 3 - figure supplement 1.**
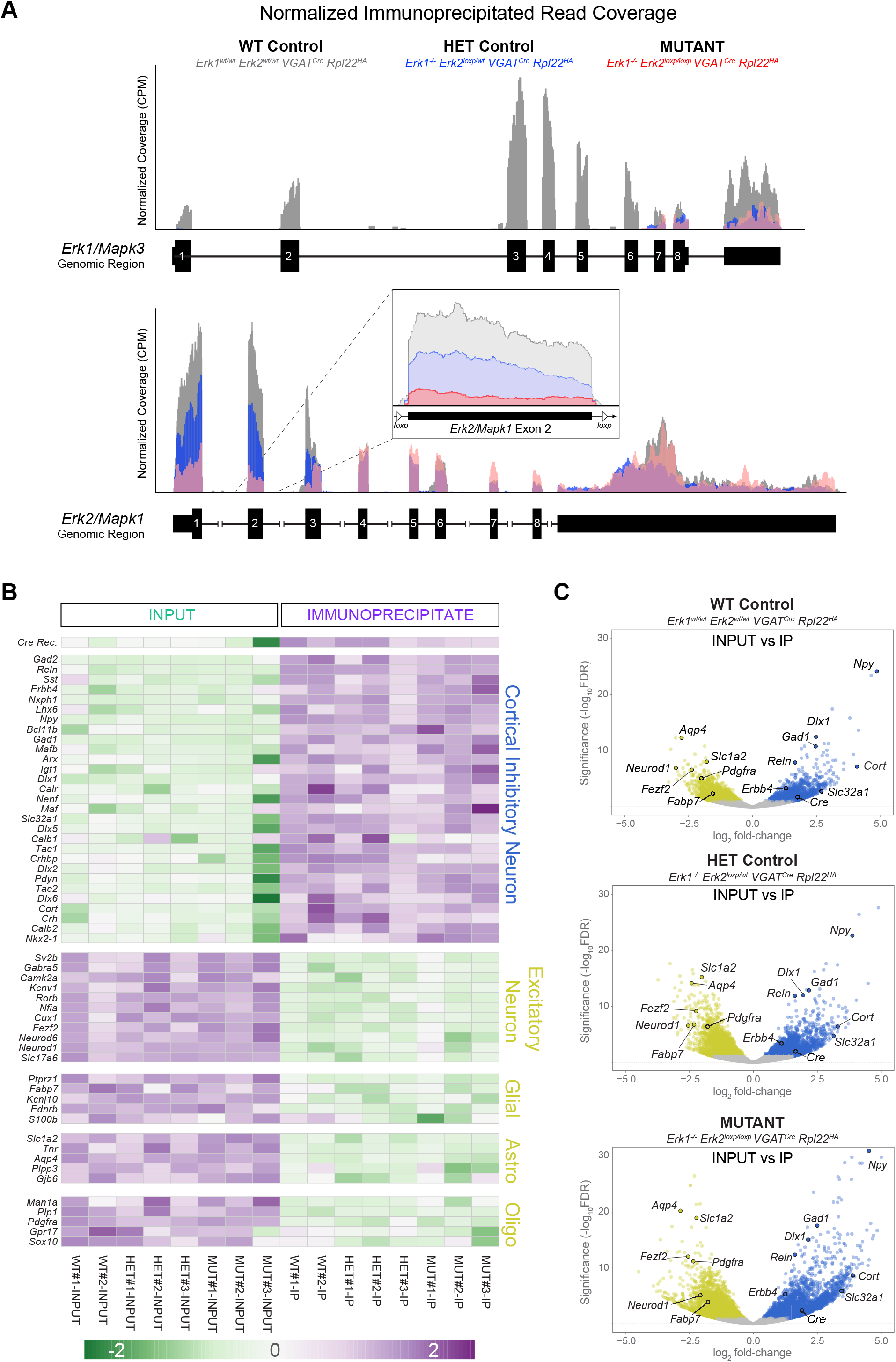
RiboTag in *Erk1/2 VGAT^Cre^* mice enriches for CIN mRNA. **(A)** Read coverage maps for *Erk1/Mapk3* and *Erk2/Mapk1* in a representative immunoprecipitated RNA sample from a wildtype (grey), heterozygous control (blue), and mutant *Erk1 Erk2 VGAT^Cre^ Rpl22^HA^* (red) cortex. As expected, *Erk1* exons 1-6 were not detected in heterozygous (N=3) and mutant (N=3) fractions relative to wildtype (N=2) samples (DEXSeq: *p*<0.005). Analysis of *Erk2/Mapk1* exon 2 (inset), which is *loxp*-flanked in conditional knockouts, revealed a statistically significant reduction in immunoprecipitated mutant samples when compared to heterozygote controls (DEXSeq: 2.5-fold reduction, *p*<0.005, N=3). **(B)** Heatmap of DeSeq2-derived normalized gene expression values for select protein coding genes with known specificity for CINs, excitatory neurons, pan-glia, astrocytes, and oligodendrocytes in all samples. Values were unit-variance scaled for each gene within a row for visualization purposes; purple values indicate relative enrichment while green values indicate relative depletion. **(C)** Volcano plots including >11,000 expressed protein coding genes in IP vs INPUT fractions within each of the three test conditions. Grey dots indicate genes not significantly changed, yellow-green dots are significantly enriched genes in INPUT fractions, while blue dots are significantly enriched genes in IP fractions. Select genes known to be expressed in distinct cell types noted in (B) are highlighted.

**Figure 4 - figure supplement 1.**
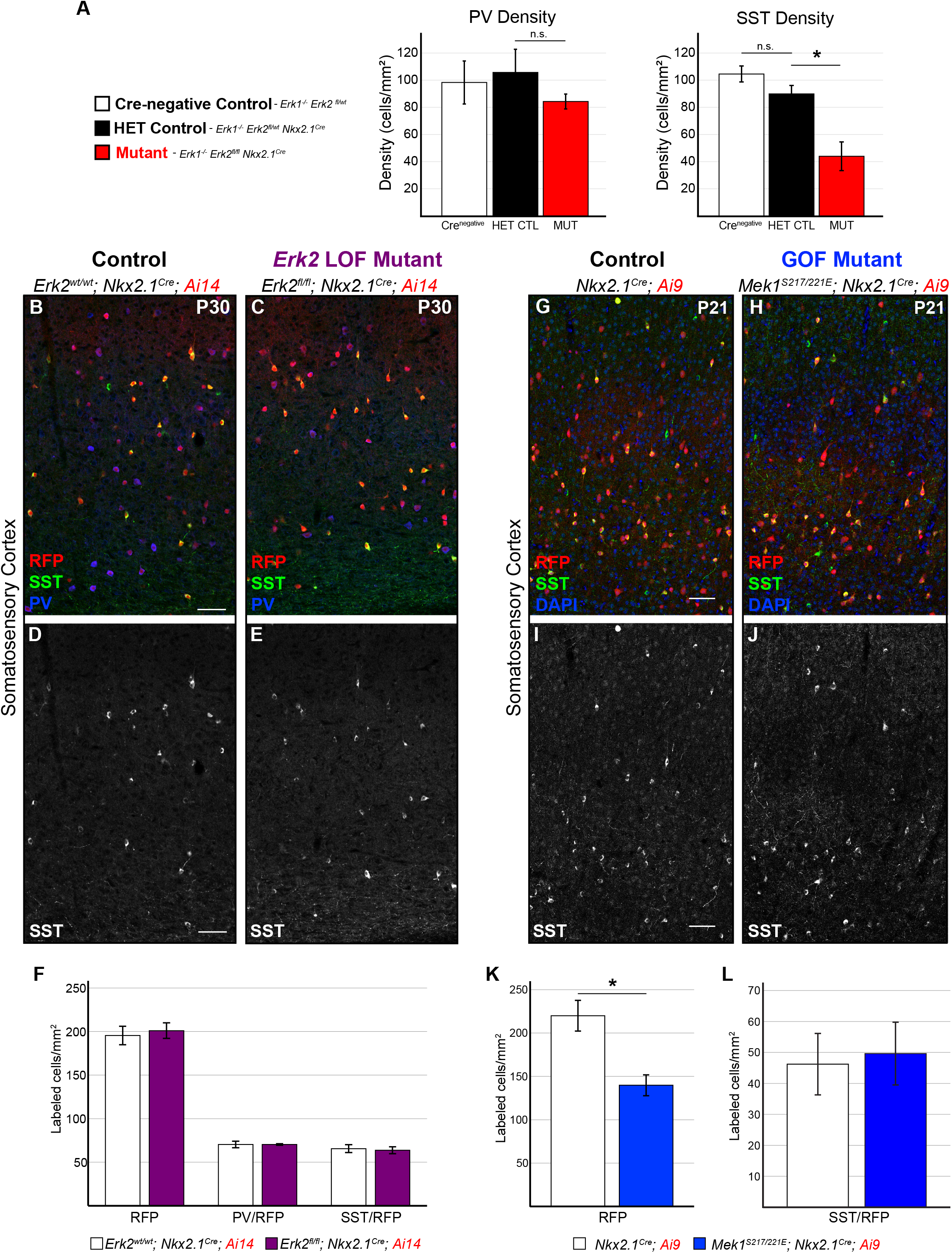
Complete loss of ERK1/2 signaling is required to alter the number of SST-expressing cells. (**A**) Quantification of total PV-expressing and SST-expressing cell density in the P60 somatosensory cortex of Cre-negative controls (*Erk1^-/-^*), HET controls (*Erk1^-/-^*; *Erk2^fl/wt^*; *Nkx2.1^Cre^*) and mutant (*Erk1^-/-^*; *Erk2^fl/fl^*; *Nkx2.1^Cre^*) mice (N=3, mean +/- SEM, one-way ANOVA followed by Fisher’s LSD; PV-*F_2,6_*=0.621, *p*>.05; SST*-F_2,6_*=16.206, *p*=.004). (**B-F**) Representative images showing CIN subtypes in the P30 primary somatosensory cortex. *Nkx2.1^Cre^*mediated deletion of *Erk2* did not alter total Cre-expressing CIN number or the density of SST-or PV-expressing CINs (quantification in **F**, N=3, mean +/- SEM, students t-test *p* >.05) (scale bar = 50 µm). **(G-L)** Hyperactivation of ERK1/2 in *Mek1^S217/221E^; Nkx2.1^Cre^* mice reduced total Cre-expressing CIN density (quantification in **K**; N=3, mean +/- SEM, students t-test *p* <.05) but did not alter SST^+^/RFP^+^-CIN density (quantification in **L**; N=3, mean +/- SEM, students t-test *p* >.05) (scale bar = 50 µm).

**Figure 5 - figure supplement 1.**
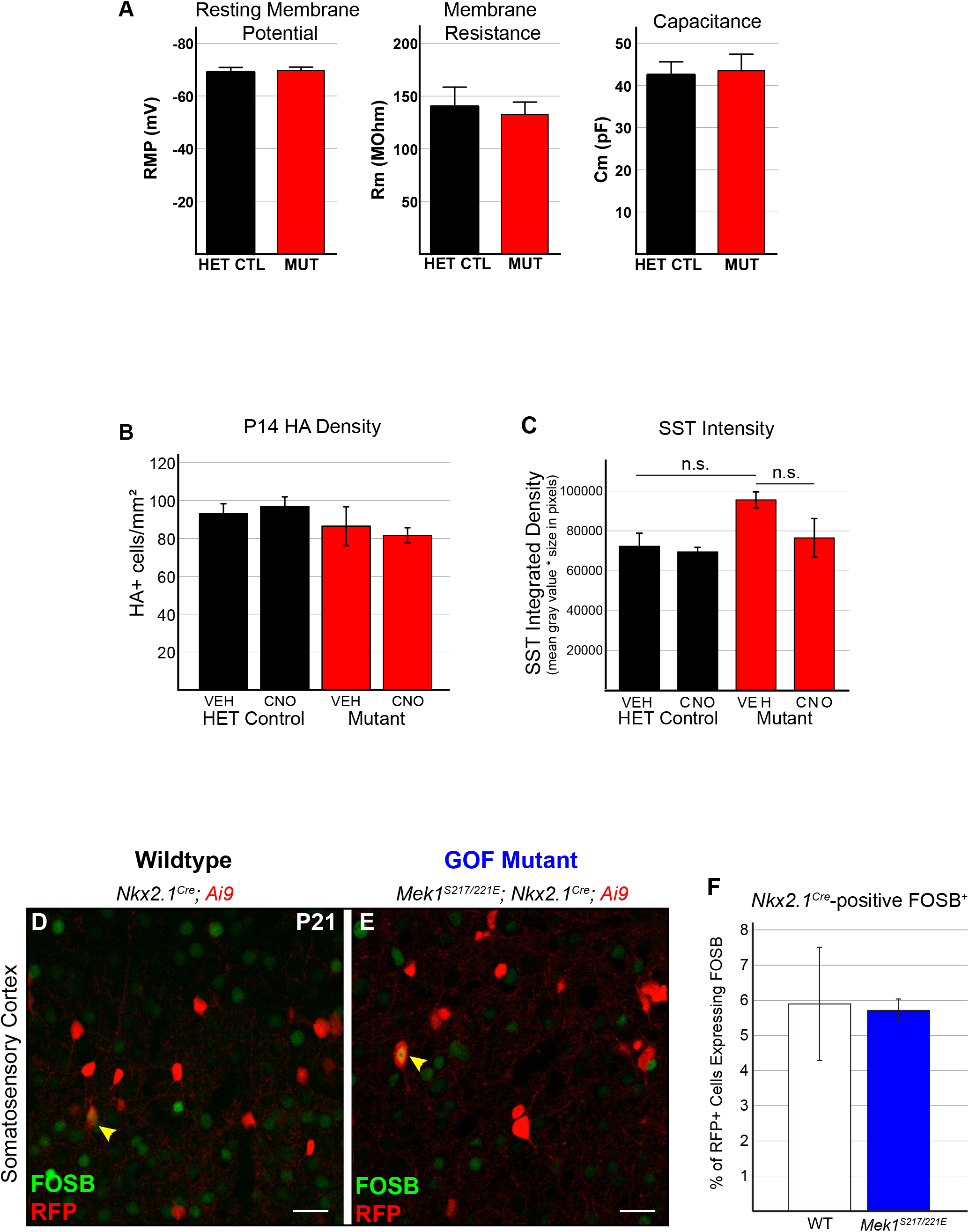
Intrinsic excitability of FS-CINs is not changed following ERK1/2 modulation. **(A)** Key FS-CIN intrinsic properties appear to be unaffected by ERK1/2 deletion, as there was no difference in resting membrane potential (*p*=0.73), membrane resistance (*p*= 0.74), or capacitance (*p*=0.85) (n=8 neurons/genotype, student’s t-test). **(B)** HA^+^ cell density was unchanged by 7 days of CNO treatment (N=4-5, one-way ANOVA, *F*_3,24_=0.62, *p*>.05). (**C**) CNO did not alter the intensity of SST immunoreactivity across genotypes (N=3 animals, minimum 12 cells/animal, one-way ANOVA, *F*_3,8_=3.446, *p*=.072). **(D-F)** Confocal images showing low expression of FOSB in P21 wildtype CINs under basal conditions (D). FOSB expression was relatively unchanged in *Mek1^S217/221E^; Nkx2.1^Cre^* CINs (E) (Quantification in **F**, mean +/- SEM, N=3, student’s t-test, *p*>.05) (scale bar = 25 µm).

## References

Adler, A., Zhao, R., Shin, M. E., Yasuda, R., & Gan, W.-B. (2019). Somatostatin-Expressing Interneurons Enable and Maintain Learning-Dependent Sequential Activation of Pyramidal Neurons. Neuron, 102(1), 202–216.e7. https://doi.org/10.1016/j.neuron.2019.01.036

Alessi, D. R., Saito, Y., Campbell, D. G., Cohen, P., Sithanandam, G., Rapp, U., Ashworth, A., Marshall, C. J., & Cowley, S. (1994). Identification of the sites in MAP kinase kinase-1 phosphorylated by p74raf-1. The EMBO Journal, 13(7), 1610–1619.

Anders, S., Reyes, A., & Huber, W. (2012). Detecting differential usage of exons from RNA-seq data. Genome Research, 22(10), 2008–2017. https://doi.org/10.1101/gr.133744.111

Anderson, T. R., Huguenard, J. R., & Prince, D. A. (2010). Differential effects of Na+-K+ ATPase blockade on cortical layer V neurons. The Journal of Physiology, 588(Pt 22), 4401–4414. https://doi.org/10.1113/jphysiol.2010.191858

Angara, K., Pai, E. L.-L., Bilinovich, S. M., Stafford, A. M., Nguyen, J. T., Li, K. X., Paul, A., Rubenstein, J. L., & Vogt, D. (2020). Nf1 deletion results in depletion of the Lhx6 transcription factor and a specific loss of parvalbumin+ cortical interneurons. Proceedings of the National Academy of Sciences, 117(11), 6189–6195. https://doi.org/10.1073/pnas.1915458117

Assimacopoulos, S., Grove, E. A., & Ragsdale, C. W. (2003). Identification of a *Pax6*-Dependent Epidermal Growth Factor Family Signaling Source at the Lateral Edge of the Embryonic Cerebral Cortex. The Journal of Neuroscience, 23(16), 6399–6403. https://doi.org/10.1523/JNEUROSCI.23-16-06399.2003

Banks, W. A., Schally, A. V., Barrera, C. M., Fasold, M. B., Durham, D. A., Csernus, V. J., Groot, K., & Kastin, A. J. (1990). Permeability of the murine blood-brain barrier to some octapeptide analogs of somatostatin. Proceedings of the National Academy of Sciences, 87(17), 6762–6766. https://doi.org/10.1073/pnas.87.17.6762

Baraban, S. C., & Tallent, M. K. (2004). Interneuron Diversity series: Interneuronal neuropeptides – endogenous regulators of neuronal excitability. Trends in Neurosciences, 27(3), 135–142. https://doi.org/10.1016/j.tins.2004.01.008

Batista-Brito, R., Machold, R., Klein, C., & Fishell, G. (2008). Gene Expression in Cortical Interneuron Precursors is Prescient of their Mature Function. Cerebral Cortex, 18(10), 2306–2317. https://doi.org/10.1093/cercor/bhm258

Ben-Ari, Y., Gaiarsa, J.-L., Tyzio, R., & Khazipov, R. (2007). GABA: A Pioneer Transmitter That Excites Immature Neurons and Generates Primitive Oscillations. Physiological Reviews, 87(4), 1215–1284. https://doi.org/10.1152/physrev.00017.2006

Ben-Shachar, S., Ou, Z., Shaw, C. A., Belmont, J. W., Patel, M. S., Hummel, M., Amato, S., Tartaglia, N., Berg, J., Sutton, V. R., Lalani, S. R., Chinault, A. C., Cheung, S. W., Lupski, J. R., & Patel, A. (2008). 22q11.2 Distal Deletion: A Recurrent Genomic Disorder Distinct from DiGeorge Syndrome and Velocardiofacial Syndrome. The American Journal of Human Genetics, 82(1), 214–221. https://doi.org/10.1016/j.ajhg.2007.09.014

Bitzenhofer, S. H., Pöpplau, J. A., Chini, M., Marquardt, A., & Hanganu-Opatz, I. L. (2021). A transient developmental increase in prefrontal activity alters network maturation and causes cognitive dysfunction in adult mice. Neuron, 109(8), 1350–1364.e6. https://doi.org/10.1016/j.neuron.2021.02.011

Blizinsky, K. D., Diaz-Castro, B., Forrest, M. P., Schürmann, B., Bach, A. P., Martin-de-Saavedra, M. D., Wang, L., Csernansky, J. G., Duan, J., & Penzes, P. (2016). Reversal of dendritic phenotypes in 16p11.2 microduplication mouse model neurons by pharmacological targeting of a network hub. Proceedings of the National Academy of Sciences, 113(30), 8520–8525. https://doi.org/10.1073/pnas.1607014113

Bomkamp, C., Tripathy, S. J., Bengtsson Gonzales, C., Hjerling-Leffler, J., Craig, A. M., & Pavlidis, P. (2019). Transcriptomic correlates of electrophysiological and morphological diversity within and across excitatory and inhibitory neuron classes. PLOS Computational Biology, 15(6), e1007113. https://doi.org/10.1371/journal.pcbi.1007113

Breunig, J. J., Levy, R., Antonuk, C. D., Molina, J., Dutra-Clarke, M., Park, H., Akhtar, A. A., Kim, G. B., Town, T., Hu, X., Bannykh, S. I., Verhaak, R. G. W., & Danielpour, M. (2015). Ets Factors Regulate Neural Stem Cell Depletion and Gliogenesis in Ras Pathway Glioma. Cell Reports, 12(2), 258–271. https://doi.org/10.1016/j.celrep.2015.06.012

Bueno, O. F. (2000). The MEK1-ERK1/2 signaling pathway promotes compensated cardiac hypertrophy in transgenic mice. The EMBO Journal, 19(23), 6341–6350. https://doi.org/10.1093/emboj/19.23.6341

Cancedda, L., Fiumelli, H., Chen, K., & Poo, M.-m. (2007). Excitatory GABA Action Is Essential for Morphological Maturation of Cortical Neurons In Vivo. Journal of Neuroscience, 27(19), 5224– 5235. https://doi.org/10.1523/JNEUROSCI.5169-06.2007

Chang, Y. S., Owen, J. P., Pojman, N. J., Thieu, T., Bukshpun, P., Wakahiro, M. L. J., Marco, E. J., Berman, J. I., Spiro, J. E., Chung, W. K., Buckner, R. L., Roberts, T. P. L., Nagarajan, S. S., Sherr, E. H., & Mukherjee, P. (2016). Reciprocal white matter alterations due to 16p11.2 chromosomal deletions versus duplications. Human Brain Mapping, 37(8), 2833–2848. https://doi.org/10.1002/hbm.23211

Chen, K., Yang, G., So, K.-F., & Zhang, L. (2019). Activation of Cortical Somatostatin Interneurons Rescues Synapse Loss and Motor Deficits after Acute MPTP Infusion. IScience, 17, 230–241. https://doi.org/10.1016/j.isci.2019.06.040

Choveau, F. S., Abderemane-Ali, F., Coyan, F. C., Es-Salah-Lamoureux, Z., Baró, I., & Loussouarn, G. (2012). Opposite Effects of the S4–S5 Linker and PIP2 on Voltage-Gated Channel Function: KCNQ1/KCNE1 and Other Channels. Frontiers in Pharmacology, 3. https://doi.org/10.3389/fphar.2012.00125

Close, J., Xu, H., De Marco Garcia, N., Batista-Brito, R., Rossignol, E., Rudy, B., & Fishell, G. (2012). Satb1 Is an Activity-Modulated Transcription Factor Required for the Terminal Differentiation and Connectivity of Medial Ganglionic Eminence-Derived Cortical Interneurons. Journal of Neuroscience, 32(49), 17690–17705. https://doi.org/10.1523/JNEUROSCI.3583-12.2012

Cohen, S. M., Ma, H., Kuchibhotla, K. V., Watson, B. O., Buzsáki, G., Froemke, R. C., & Tsien, R. W. (2016). Excitation-Transcription Coupling in Parvalbumin-Positive Interneurons Employs a Novel CaM Kinase-Dependent Pathway Distinct from Excitatory Neurons. Neuron, 90(2), 292–307. https://doi.org/10.1016/j.neuron.2016.03.001

Courchesne, E., Gazestani, V. H., & Lewis, N. E. (2020). Prenatal Origins of ASD: The When, What, and How of ASD Development. Trends in Neurosciences, 43(5), 326–342. https://doi.org/10.1016/j.tins.2020.03.005

Cowley, S. (1994). Activation of MAP kinase kinase is necessary and sufficient for PC12 differentiation and for transformation of NIH 3T3 cells. Cell, 77(6), 841–852. https://doi.org/10.1016/0092-8674(94)90133-3

Cristo, G. D., Berardi, N., Cancedda, L., Pizzorusso, T., Putignano, E., Ratto, G. M., & Maffei, L. (2001). Requirement of ERK Activation for Visual Cortical Plasticity. Science, 292(5525), 2337–2340. https://doi.org/10.1126/science.1059075

Cummings, K. A., & Clem, R. L. (2020). Prefrontal somatostatin interneurons encode fear memory. Nature Neuroscience, 23(1), 61–74. https://doi.org/10.1038/s41593-019-0552-7

Davies, P., Katzman, R., & Terry, R. D. (1980). Reduced somatostatin-like immunoreactivity in cerebral cortex from cases of Alzheimer disease and Alzheimer senile dementa. Nature, 288(5788), 279–280. https://doi.org/10.1038/288279a0

De Marco García, N. V., Karayannis, T., & Fishell, G. (2011). Neuronal activity is required for the development of specific cortical interneuron subtypes. Nature, 472(7343), 351–355. https://doi.org/10.1038/nature09865

DeFelipe, J., López-Cruz, P. L., Benavides-Piccione, R., Bielza, C., Larrañaga, P., Anderson, S., Burkhalter, A., Cauli, B., Fairén, A., Feldmeyer, D., Fishell, G., Fitzpatrick, D., Freund, T. F., González-Burgos, G., Hestrin, S., Hill, S., Hof, P. R., Huang, J., Jones, E. G., … Ascoli, G. A. (2013). New insights into the classification and nomenclature of cortical GABAergic interneurons. Nature Reviews Neuroscience, 14(3), 202–216. https://doi.org/10.1038/nrn3444

Dehorter, N., Ciceri, G., Bartolini, G., Lim, L., del Pino, I., & Marín, O. (2015). Tuning of fast-spiking interneuron properties by an activity-dependent transcriptional switch. Science, 349(6253), 1216– 1220. https://doi.org/10.1126/science.aab3415

del Rio, J., de Lecea, L., Ferrer, I., & Soriano, E. (1994). The development of parvalbumin-immunoreactivity in the neocortex of the mouse. Developmental Brain Research, 81(2), 247–259. https://doi.org/10.1016/0165-3806(94)90311-5

Denaxa, M., Kalaitzidou, M., Garefalaki, A., Achimastou, A., Lasrado, R., Maes, T., & Pachnis, V. (2012). Maturation-Promoting Activity of SATB1 in MGE-Derived Cortical Interneurons. Cell Reports, 2(5), 1351–1362. https://doi.org/10.1016/j.celrep.2012.10.003

Denaxa, M., Neves, G., Rabinowitz, A., Kemlo, S., Liodis, P., Burrone, J., & Pachnis, V. (2018). Modulation of Apoptosis Controls Inhibitory Interneuron Number in the Cortex. Cell Reports, 22(7), 1710–1721. https://doi.org/10.1016/j.celrep.2018.01.064

Dinh Duong, T. A., Hoshiba, Y., Saito, K., Kawasaki, K., Ichikawa, Y., Matsumoto, N., Shinmyo, Y., & Kawasaki, H. (2019). FGF Signaling Directs the Cell Fate Switch from Neurons to Astrocytes in the Developing Mouse Cerebral Cortex. The Journal of Neuroscience, 39(31), 6081–6094. https://doi.org/10.1523/JNEUROSCI.2195-18.2019

Dobin, A., Davis, C. A., Schlesinger, F., Drenkow, J., Zaleski, C., Jha, S., Batut, P., Chaisson, M., & Gingeras, T. R. (2013). STAR: Ultrafast universal RNA-seq aligner. *Bioinformatics (Oxford*, England), 29(1), 15–21. https://doi.org/10.1093/bioinformatics/bts635

Edgar, R., Domrachev, M., & Lash, A. E. (2002). Gene Expression Omnibus: NCBI gene expression and hybridization array data repository. Nucleic Acids Research, 30(1), 207–210. https://doi.org/10.1093/nar/30.1.207

Engin, E., & Treit, D. (2009). Anxiolytic and antidepressant actions of somatostatin: The role of sst2 and sst3 receptors. Psychopharmacology, 206(2), 281–289. https://doi.org/10.1007/s00213-009-1605-5

Exposito-Alonso, D., Osório, C., Bernard, C., Pascual-García, S., del Pino, I., Marín, O., & Rico, B. (2020). Subcellular sorting of neuregulins controls the assembly of excitatory-inhibitory cortical circuits. ELife, 9, e57000. https://doi.org/10.7554/eLife.57000

Faedo, A., Borello, U., & Rubenstein, J. L. R. (2010). Repression of Fgf Signaling by Sprouty1-2 Regulates Cortical Patterning in Two Distinct Regions and Times. Journal of Neuroscience, 30(11), 4015–4023. https://doi.org/10.1523/JNEUROSCI.0307-10.2010

Fattah, M., Raman, M. M., Reiss, A. L., & Green, T. (2021). PTPN11 Mutations in the Ras-MAPK Signaling Pathway Affect Human White Matter Microstructure. Cerebral Cortex, 31(3), 1489– 1499. https://doi.org/10.1093/cercor/bhaa299

Fazzari, P., Paternain, A. V., Valiente, M., Pla, R., Luján, R., Lloyd, K., Lerma, J., Marín, O., & Rico, B. (2010). Control of cortical GABA circuitry development by Nrg1 and ErbB4 signalling. Nature, 464(7293), 1376–1380. https://doi.org/10.1038/nature08928

Fee, C., Banasr, M., & Sibille, E. (2017). Somatostatin-Positive Gamma-Aminobutyric Acid Interneuron Deficits in Depression: Cortical Microcircuit and Therapeutic Perspectives. Biological Psychiatry, 82(8), 549–559. https://doi.org/10.1016/j.biopsych.2017.05.024

Fernandez, B. A., Roberts, W., Chung, B., Weksberg, R., Meyn, S., Szatmari, P., Joseph-George, A. M., MacKay, S., Whitten, K., Noble, B., Vardy, C., Crosbie, V., Luscombe, S., Tucker, E., Turner, L., Marshall, C. R., & Scherer, S. W. (2010). Phenotypic spectrum associated with de novo and inherited deletions and duplications at 16p11.2 in individuals ascertained for diagnosis of autism spectrum disorder. Journal of Medical Genetics, 47(3), 195–203. https://doi.org/10.1136/jmg.2009.069369

Filges, I., Sparagana, S., Sargent, M., Selby, K., Schlade-Bartusiak, K., Lueder, G. T., Robichaux-Viehoever, A., Schlaggar, B. L., Shimony, J. S., & Shinawi, M. (2014). Brain MRI abnormalities and spectrum of neurological and clinical findings in three patients with proximal 16p11.2 microduplication. American Journal of Medical Genetics. Part A, 164A(8), 2003–2012. https://doi.org/10.1002/ajmg.a.36605

Forloni, G., Hohmann, C., & Coyle, J. T. (1990). Developmental expression of somatostatin in mouse brain. I. Immunocytochemical studies. Developmental Brain Research, 53(1), 6–25. https://doi.org/10.1016/0165-3806(90)90120-N

Fu, Y., Tucciarone, J. M., Espinosa, J. S., Sheng, N., Darcy, D. P., Nicoll, R. A., Huang, Z. J., & Stryker, M. P. (2014). A cortical circuit for gain control by behavioral state. Cell, 156(6), 1139–1152. https://doi.org/10.1016/j.cell.2014.01.050

Furusho, M., Ishii, A., & Bansal, R. (2017). Signaling by FGF Receptor 2, Not FGF Receptor 1, Regulates Myelin Thickness through Activation of ERK1/2-MAPK, Which Promotes mTORC1 Activity in an Akt-Independent Manner. The Journal of Neuroscience: The Official Journal of the Society for Neuroscience, 37(11), 2931–2946. https://doi.org/10.1523/JNEUROSCI.3316-16.2017

Furusho, M., Kaga, Y., Ishii, A., Hébert, J. M., & Bansal, R. (2011). Fibroblast growth factor signaling is required for the generation of oligodendrocyte progenitors from the embryonic forebrain. The Journal of Neuroscience: The Official Journal of the Society for Neuroscience, 31(13), 5055– 5066. https://doi.org/10.1523/JNEUROSCI.4800-10.2011

Fyffe-Maricich, S. L., Karlo, J. C., Landreth, G. E., & Miller, R. H. (2011). The ERK2 Mitogen-Activated Protein Kinase Regulates the Timing of Oligodendrocyte Differentiation. Journal of Neuroscience, 31(3), 843–850. https://doi.org/10.1523/JNEUROSCI.3239-10.2011

Glorioso, C., Sabatini, M., Unger, T., Hashimoto, T., Monteggia, L. M., Lewis, D. A., & Mirnics, K. (2006). Specificity and timing of neocortical transcriptome changes in response to BDNF gene ablation during embryogenesis or adulthood. Molecular Psychiatry, 11(7), 633–648. https://doi.org/10.1038/sj.mp.4001835

Goddeyne, C., Nichols, J., Wu, C., & Anderson, T. (2015). Repetitive mild traumatic brain injury induces ventriculomegaly and cortical thinning in juvenile rats. Journal of Neurophysiology, 113(9), 3268–3280. https://doi.org/10.1152/jn.00970.2014

Gour, A., Boergens, K. M., Heike, N., Hua, Y., Laserstein, P., Song, K., & Helmstaedter, M. (2021). Postnatal connectomic development of inhibition in mouse barrel cortex. Science, 371(6528), eabb4534. https://doi.org/10.1126/science.abb4534

Grilli, M., Raiteri, L., & Pittaluga, A. (2004). Somatostatin inhibits glutamate release from mouse cerebrocortical nerve endings through presynaptic sst2 receptors linked to the adenylyl cyclase– protein kinase A pathway. Neuropharmacology, 46(3), 388–396. https://doi.org/10.1016/j.neuropharm.2003.09.012

Guenthner, C. J., Miyamichi, K., Yang, H. H., Heller, H. C., & Luo, L. (2013). Permanent Genetic Access to Transiently Active Neurons via TRAP: Targeted Recombination in Active Populations. Neuron, 78(5), 773–784. https://doi.org/10.1016/j.neuron.2013.03.025

Gutmann, D. H., Wu, Y. L., Hedrick, N. M., Zhu, Y., Guha, A., & Parada, L. F. (2001). Heterozygosity for the neurofibromatosis 1 (NF1) tumor suppressor results in abnormalities in cell attachment, spreading and motility in astrocytes. Human Molecular Genetics, 10(26), 3009–3016. https://doi.org/10.1093/hmg/10.26.3009

Hanson, E., Armbruster, M., Lau, L. A., Sommer, M. E., Klaft, Z.-J., Swanger, S. A., Traynelis, S. F., Moss, S. J., Noubary, F., Chadchankar, J., & Dulla, C. G. (2019). Tonic Activation of GluN2C/GluN2D-Containing NMDA Receptors by Ambient Glutamate Facilitates Cortical Interneuron Maturation. The Journal of Neuroscience: The Official Journal of the Society for Neuroscience, 39(19), 3611–3626. https://doi.org/10.1523/JNEUROSCI.1392-18.2019

Heavner, W. E., & Smith, S. E. P. (2020). Resolving the Synaptic versus Developmental Dichotomy of Autism Risk Genes. Trends in Neurosciences, 43(4), 227–241. https://doi.org/10.1016/j.tins.2020.01.009

Holter, M. C., Hewitt, L. T., Nishimura, K. J., Knowles, S. J., Bjorklund, G. R., Shah, S., Fry, N. R., Rees, K. P., Gupta, T. A., Daniels, C. W., Li, G., Marsh, S., Treiman, D. M., Olive, M. F., Anderson, T. R., Sanabria, F., Snider, W. D., & Newbern, J. M. (2021). Hyperactive MEK1 Signaling in Cortical GABAergic Neurons Promotes Embryonic Parvalbumin Neuron Loss and Defects in Behavioral Inhibition. *Cerebral Cortex*, bhaa413. https://doi.org/10.1093/cercor/bhaa413

Holter, M. C., Hewitt, Lauren. T., Koebele, S. V., Judd, J. M., Xing, L., Bimonte-Nelson, H. A., Conrad, C. D., Araki, T., Neel, B. G., Snider, W. D., & Newbern, J.M. (2019). The Noonan Syndrome-linked Raf1L613V mutation drives increased glial number in the mouse cortex and enhanced learning. PLOS Genetics, 15(4), e1008108. https://doi.org/10.1371/journal.pgen.1008108

Hong, E. J., McCord, A. E., & Greenberg, M. E. (2008). A Biological Function for the Neuronal Activity-Dependent Component of Bdnf Transcription in the Development of Cortical Inhibition. Neuron, 60(4), 610–624. https://doi.org/10.1016/j.neuron.2008.09.024

Hrvatin, S., Hochbaum, D. R., Nagy, M. A., Cicconet, M., Robertson, K., Cheadle, L., Zilionis, R., Ratner, A., Borges-Monroy, R., Klein, A. M., Sabatini, B. L., & Greenberg, M. E. (2018). Single-cell analysis of experience-dependent transcriptomic states in the mouse visual cortex. Nature Neuroscience, 21(1), 120–129. https://doi.org/10.1038/s41593-017-0029-5

Hu, J. S., Vogt, D., Sandberg, M., & Rubenstein, J. L. (2017). Cortical interneuron development: A tale of time and space. Development, 144(21), 3867–3878. https://doi.org/10.1242/dev.132852

Huang, Z. J., Kirkwood, A., Pizzorusso, T., Porciatti, V., Morales, B., Bear, M. F., Maffei, L., & Tonegawa, S. (1999). BDNF Regulates the Maturation of Inhibition and the Critical Period of Plasticity in Mouse Visual Cortex. Cell, 98(6), 739–755. https://doi.org/10.1016/S0092-8674(00)81509-3

Hutton, S. R., Otis, J. M., Kim, E. M., Lamsal, Y., Stuber, G. D., & Snider, W. D. (2017). ERK/MAPK Signaling Is Required for Pathway-Specific Striatal Motor Functions. The Journal of Neuroscience: The Official Journal of the Society for Neuroscience, 37(34), 8102–8115. https://doi.org/10.1523/JNEUROSCI.0473-17.2017

Imamura, O., Pagès, G., Pouysségur, J., Endo, S., & Takishima, K. (2010). ERK1 and ERK2 are required for radial glial maintenance and cortical lamination. Genes to Cells: Devoted to Molecular & Cellular Mechanisms, 15(10), 1072–1088. https://doi.org/10.1111/j.1365-2443.2010.01444.x

Ishii, A., Furusho, M., Dupree, J. L., & Bansal, R. (2016). Strength of ERK1/2 MAPK Activation Determines Its Effect on Myelin and Axonal Integrity in the Adult CNS. The Journal of Neuroscience: The Official Journal of the Society for Neuroscience, 36(24), 6471–6487. https://doi.org/10.1523/JNEUROSCI.0299-16.2016

Ishii, A., Fyffe-Maricich, S. L., Furusho, M., Miller, R. H., & Bansal, R. (2012). ERK1/ERK2 MAPK Signaling is Required to Increase Myelin Thickness Independent of Oligodendrocyte Differentiation and Initiation of Myelination. Journal of Neuroscience, 32(26), 8855–8864. https://doi.org/10.1523/JNEUROSCI.0137-12.2012

Itami, C., Kimura, F., & Nakamura, S. (2007). Brain-Derived Neurotrophic Factor Regulates the Maturation of Layer 4 Fast-Spiking Cells after the Second Postnatal Week in the Developing Barrel Cortex. Journal of Neuroscience, 27(9), 2241–2252. https://doi.org/10.1523/JNEUROSCI.3345-06.2007

Jendryka, M., Palchaudhuri, M., Ursu, D., van der Veen, B., Liss, B., Kätzel, D., Nissen, W., & Pekcec, A. (2019). Pharmacokinetic and pharmacodynamic actions of clozapine-N-oxide, clozapine, and compound 21 in DREADD-based chemogenetics in mice. Scientific Reports, 9(1), 4522. https://doi.org/10.1038/s41598-019-41088-2

Kaneko, K., Currin, C. B., Goff, K. M., Wengert, E. R., Somarowthu, A., Vogels, T. P., & Goldberg, E. M. (2022). Developmentally regulated impairment of parvalbumin interneuron synaptic transmission in an experimental model of Dravet syndrome. Cell Reports, 38(13), 110580. https://doi.org/10.1016/j.celrep.2022.110580

Kawaguchi, Y. (1997). GABAergic cell subtypes and their synaptic connections in rat frontal cortex. Cerebral Cortex, 7(6), 476–486. https://doi.org/10.1093/cercor/7.6.476

Kessaris, N., Fogarty, M., Iannarelli, P., Grist, M., Wegner, M., & Richardson, W. D. (2006). Competing waves of oligodendrocytes in the forebrain and postnatal elimination of an embryonic lineage. Nature Neuroscience, 9(2), 173–179. https://doi.org/10.1038/nn1620

Kessaris, N., Jamen, F., Rubin, L. L., & Richardson, W. D. (2004). Cooperation between sonic hedgehog and fibroblast growth factor/MAPK signalling pathways in neocortical precursors. *Development (Cambridge*, England), 131(6), 1289–1298. https://doi.org/10.1242/dev.01027

Kiviniemi, A., Gardberg, M., Frantzén, J., Pesola, M., Vuorinen, V., Parkkola, R., Tolvanen, T., Suilamo, S., Johansson, J., Luoto, P., Kemppainen, J., Roivainen, A., & Minn, H. (2015). Somatostatin receptor subtype 2 in high-grade gliomas: PET/CT with 68Ga-DOTA-peptides, correlation to prognostic markers, and implications for targeted radiotherapy. EJNMMI Research, 5(1), 25. https://doi.org/10.1186/s13550-015-0106-2

Klemke, R. L., Cai, S., Giannini, A. L., Gallagher, P. J., de Lanerolle, P., & Cheresh, D. A. (1997). Regulation of cell motility by mitogen-activated protein kinase. The Journal of Cell Biology, 137(2), 481–492. https://doi.org/10.1083/jcb.137.2.481

Klesse, L. J., Meyers, K. A., Marshall, C. J., & Parada, L. F. (1999). Nerve growth factor induces survival and differentiation through two distinct signaling cascades in PC12 cells. Oncogene, 18(12), 2055–2068. https://doi.org/10.1038/sj.onc.1202524

Kluge, C., Stoppel, C., Szinyei, C., Stork, O., & Pape, H.-C. (2008). Role of the somatostatin system in contextual fear memory and hippocampal synaptic plasticity. Learning & Memory, 15(4), 252–260. https://doi.org/10.1101/lm.793008

Knowles, S. J., Stafford, A. M., Zaman, T., Angara, K., Williams, M. R., Newbern, J. M., & Vogt, D. (2022). *Distinct hyperactive RAS/MAPK alleles converge on common GABAergic interneuron core programs* [Preprint]. Neuroscience. https://doi.org/10.1101/2022.08.04.502867

Krencik, R., Hokanson, K. C., Narayan, A. R., Dvornik, J., Rooney, G. E., Rauen, K. A., Weiss, L. A., Rowitch, D. H., & Ullian, E. M. (2015). Dysregulation of astrocyte extracellular signaling in Costello syndrome. Science Translational Medicine, 7(286), 286ra66. https://doi.org/10.1126/scitranslmed.aaa5645

Krenz, M., Gulick, J., Osinska, H. E., Colbert, M. C., Molkentin, J. D., & Robbins, J. (2008). Role of ERK1/2 signaling in congenital valve malformations in Noonan syndrome. Proceedings of the National Academy of Sciences, 105(48), 18930–18935. https://doi.org/10.1073/pnas.0806556105

Lajiness, J. D., Snider, P., Wang, J., Feng, G.-S., Krenz, M., & Conway, S. J. (2014). SHP-2 deletion in postmigratory neural crest cells results in impaired cardiac sympathetic innervation. Proceedings of the National Academy of Sciences, 111(14), E1374–E1382. https://doi.org/10.1073/pnas.1319208111

Lavoie, H., Gagnon, J., & Therrien, M. (2020). ERK signalling: A master regulator of cell behaviour, life and fate. Nature Reviews Molecular Cell Biology, 21(10), 607–632. https://doi.org/10.1038/s41580-020-0255-7

Lee, C., Harkin, E. F., Yin, X., Naud, R., & Chen, S. (2022). Cell-type-specific responses to associative learning in the primary motor cortex. ELife, 11, e72549. https://doi.org/10.7554/eLife.72549

Lee, D. R., Rhodes, C., Mitra, A., Zhang, Y., Maric, D., Dale, R. K., & Petros, T. J. (2022). Transcriptional heterogeneity of ventricular zone cells in the ganglionic eminences of the mouse forebrain. ELife, 11, e71864. https://doi.org/10.7554/eLife.71864

Li, S., Mattar, P., Dixit, R., Lawn, S. O., Wilkinson, G., Kinch, C., Eisenstat, D., Kurrasch, D. M., Chan, J. A., & Schuurmans, C. (2014). RAS/ERK Signaling Controls Proneural Genetic Programs in Cortical Development and Gliomagenesis. Journal of Neuroscience, 34(6), 2169–2190. https://doi.org/10.1523/JNEUROSCI.4077-13.2014

Li, X., Newbern, J. M., Wu, Y., Morgan-Smith, M., Zhong, J., Charron, J., & Snider, W. D. (2012). MEK Is a Key Regulator of Gliogenesis in the Developing Brain. Neuron, 75(6), 1035–1050. https://doi.org/10.1016/j.neuron.2012.08.031

Liguz-Lecznar, M., Urban-Ciecko, J., & Kossut, M. (2016). Somatostatin and Somatostatin-Containing Neurons in Shaping Neuronal Activity and Plasticity. Frontiers in Neural Circuits, 10. https://doi.org/10.3389/fncir.2016.00048

Lim, L., Mi, D., Llorca, A., & Marín, O. (2018). Development and Functional Diversification of Cortical Interneurons. Neuron, 100(2), 294–313. https://doi.org/10.1016/j.neuron.2018.10.009

Linhares, N. D., Freire, M. C. M., Cardenas, R. G. C. do C. L., Pena, H. B., Lachlan, K., Dallapiccola, B., Bacino, C., Delobel, B., James, P., Thuresson, A.-C., Annerén, G., & Pena, S. D. J. (2016). 1p13.2 deletion displays clinical features overlapping Noonan syndrome, likely related to NRAS gene haploinsufficiency. Genetics and Molecular Biology, 39(3), 349–357. https://doi.org/10.1590/1678-4685-GMB-2016-0049

Lodato, S., Rouaux, C., Quast, K. B., Jantrachotechatchawan, C., Studer, M., Hensch, T. K., & Arlotta, P. (2011). Excitatory Projection Neuron Subtypes Control the Distribution of Local Inhibitory Interneurons in the Cerebral Cortex. Neuron, 69(4), 763–779. https://doi.org/10.1016/j.neuron.2011.01.015

López-Juárez, A., Titus, H. E., Silbak, S. H., Pressler, J. W., Rizvi, T. A., Bogard, M., Bennett, M. R., Ciraolo, G., Williams, M. T., Vorhees, C. V., & Ratner, N. (2017). Oligodendrocyte Nf1 Controls Aberrant Notch Activation and Regulates Myelin Structure and Behavior. Cell Reports, 19(3), 545–557. https://doi.org/10.1016/j.celrep.2017.03.073

Lord, C., Brugha, T. S., Charman, T., Cusack, J., Dumas, G., Frazier, T., Jones, E. J. H., Jones, R. M., Pickles, A., State, M. W., Taylor, J. L., & Veenstra-VanderWeele, J. (2020). Autism spectrum disorder. Nature Reviews Disease Primers, 6(1), 5. https://doi.org/10.1038/s41572-019-0138-4

Love, M. I., Huber, W., & Anders, S. (2014). Moderated estimation of fold change and dispersion for RNA-seq data with DESeq2. Genome Biology, 15(12), 550. https://doi.org/10.1186/s13059-014-0550-8

Ma, S., Skarica, M., Li, Q., Xu, C., Risgaard, R. D., Tebbenkamp, A. T. N., Mato-Blanco, X., Kovner, R., Krsnik, Ž., de Martin, X., Luria, V., Martí-Pérez, X., Liang, D., Karger, A., Schmidt, D. K., Gomez-Sanchez, Z., Qi, C., Gobeske, K. T., Pochareddy, S., … Sestan, N. (2022). Molecular and cellular evolution of the primate dorsolateral prefrontal cortex. Science, 377(6614), eabo7257. https://doi.org/10.1126/science.abo7257

Madisen, L., Zwingman, T. A., Sunkin, S. M., Oh, S. W., Zariwala, H. A., Gu, H., Ng, L. L., Palmiter, R. D., Hawrylycz, M. J., Jones, A. R., Lein, E. S., & Zeng, H. (2010). A robust and high-throughput Cre reporting and characterization system for the whole mouse brain. Nature Neuroscience, 13(1), 133–140. https://doi.org/10.1038/nn.2467

Magno, L., Asgarian, Z., Pendolino, V., Velona, T., Mackintosh, A., Lee, F., Stryjewska, A., Zimmer, C., Guillemot, F., Farrant, M., Clark, B., & Kessaris, N. (2021). Transient developmental imbalance of cortical interneuron subtypes presages long-term changes in behavior. Cell Reports, 35(11), 109249. https://doi.org/10.1016/j.celrep.2021.109249

Maisonpierre, P. C., Belluscio, L., Friedman, B., Alderson, R. F., Wiegand, S. J., Furth, M. E., Lindsay, R. M., & Yancopoulos, G. D. (1990). NT-3, BDNF, and NGF in the developing rat nervous system: Parallel as well as reciprocal patterns of expression. Neuron, 5(4), 501–509. https://doi.org/10.1016/0896-6273(90)90089-X

Malik, R., Pai, E. L.-L., Rubin, A. N., Stafford, A. M., Angara, K., Minasi, P., Rubenstein, J. L., Sohal, V. S., & Vogt, D. (2019). Tsc1 represses parvalbumin expression and fast-spiking properties in somatostatin lineage cortical interneurons. Nature Communications, 10(1), 4994. https://doi.org/10.1038/s41467-019-12962-4

Mardinly, A. R., Spiegel, I., Patrizi, A., Centofante, E., Bazinet, J. E., Tzeng, C. P., Mandel-Brehm, C., Harmin, D. A., Adesnik, H., Fagiolini, M., & Greenberg, M. E. (2016). Sensory experience regulates cortical inhibition by inducing IGF1 in VIP neurons. Nature, 531(7594), 371–375. https://doi.org/10.1038/nature17187

Marsh, E. D., Nasrallah, M. P., Walsh, C., Murray, K. A., Nicole Sunnen, C., McCoy, A., & Golden, J. A. (2016). Developmental interneuron subtype deficits after targeted loss of Arx. BMC Neuroscience, 17(1), 35. https://doi.org/10.1186/s12868-016-0265-8

Martin, D., Xu, J., Porretta, C., & Nichols, C. D. (2017). Neurocytometry: Flow Cytometric Sorting of Specific Neuronal Populations from Human and Rodent Brain. ACS Chemical Neuroscience, 8(2), 356–367. https://doi.org/10.1021/acschemneuro.6b00374

Matthews, R. P., Guthrie, C. R., Wailes, L. M., Zhao, X., Means, A. R., & McKnight, G. S. (1994). Calcium/calmodulin-dependent protein kinase types II and IV differentially regulate CREB-dependent gene expression. Molecular and Cellular Biology, 14(9), 6107–6116. https://doi.org/10.1128/MCB.14.9.6107

Mayer, C., Hafemeister, C., Bandler, R. C., Machold, R., Batista Brito, R., Jaglin, X., Allaway, K., Butler, A., Fishell, G., & Satija, R. (2018). Developmental diversification of cortical inhibitory interneurons. Nature, 555(7697), 457–462. https://doi.org/10.1038/nature25999

McCormick, D. A., Connors, B. W., Lighthall, J. W., & Prince, D. A. (1985). Comparative electrophysiology of pyramidal and sparsely spiny stellate neurons of the neocortex. Journal of Neurophysiology, 54(4), 782–806. https://doi.org/10.1152/jn.1985.54.4.782

McKeage, K. (2015). Pasireotide in Acromegaly: A Review. Drugs, 75(9), 1039–1048. https://doi.org/10.1007/s40265-015-0413-y

McKenzie, M. G., Cobbs, L. V., Dummer, P. D., Petros, T. J., Halford, M. M., Stacker, S. A., Zou, Y., Fishell, G. J., & Au, E. (2019). Non-canonical Wnt Signaling through Ryk Regulates the Generation of Somatostatin- and Parvalbumin-Expressing Cortical Interneurons. Neuron, 103(5), 853–864.e4. https://doi.org/10.1016/j.neuron.2019.06.003

Mi, D., Li, Z., Lim, L., Li, M., Moissidis, M., Yang, Y., Gao, T., Hu, T. X., Pratt, T., Price, D. J., Sestan, N., & Marín, O. (2018). Early emergence of cortical interneuron diversity in the mouse embryo. Science, 360(6384), 81–85. https://doi.org/10.1126/science.aar6821

Miller, M. N., Okaty, B. W., Kato, S., & Nelson, S. B. (2011). Activity-dependent changes in the firing properties of neocortical fast-spiking interneurons in the absence of large changes in gene expression. Developmental Neurobiology, 71(1), 62–70. https://doi.org/10.1002/dneu.20811

Minocha, S., Valloton, D., Ypsilanti, A. R., Fiumelli, H., Allen, E. A., Yanagawa, Y., Marin, O., Chédotal, A., Hornung, J.-P., & Lebrand, C. (2015). Nkx2.1-derived astrocytes and neurons together with Slit2 are indispensable for anterior commissure formation. Nature Communications, 6, 6887. https://doi.org/10.1038/ncomms7887

Miyoshi, G., Butt, S. J. B., Takebayashi, H., & Fishell, G. (2007). Physiologically Distinct Temporal Cohorts of Cortical Interneurons Arise from Telencephalic Olig2-Expressing Precursors. Journal of Neuroscience, 27(29), 7786–7798. https://doi.org/10.1523/JNEUROSCI.1807-07.2007

Monory, K., Massa, F., Egertová, M., Eder, M., Blaudzun, H., Westenbroek, R., Kelsch, W., Jacob, W., Marsch, R., Ekker, M., Long, J., Rubenstein, J. L., Goebbels, S., Nave, K.-A., During, M., Klugmann, M., Wölfel, B., Dodt, H.-U., Zieglgänsberger, W., … Lutz, B. (2006). The Endocannabinoid System Controls Key Epileptogenic Circuits in the Hippocampus. Neuron, 51(4), 455–466. https://doi.org/10.1016/j.neuron.2006.07.006

Montminy, M. R., & Bilezikjian, L. M. (1987). Binding of a nuclear protein to the cyclic-AMP response element of the somatostatin gene. Nature, 328(6126), 175–178. https://doi.org/10.1038/328175a0

Mori, M. X., Itsuki, K., Hase, H., Sawamura, S., Kurokawa, T., Mori, Y., & Inoue, R. (2015). Dynamics of receptor-operated Ca2+ currents through TRPC channels controlled via the PI(4,5)P2-PLC signaling pathway. Frontiers in Pharmacology, 6. https://doi.org/10.3389/fphar.2015.00022

Moyses-Oliveira, M., Yadav, R., Erdin, S., & Talkowski, M. E. (2020). New gene discoveries highlight functional convergence in autism and related neurodevelopmental disorders. Current Opinion in Genetics & Development, 65, 195–206. https://doi.org/10.1016/j.gde.2020.07.001

Munguba, H., Nikouei, K., Hochgerner, H., Oberst, P., Kouznetsova, A., Ryge, J., Batista-Brito, R., Munoz-Manchado, A. B., Close, J., Linnarsson, S., & Hjerling-Leffler, J. (2019). *Transcriptional maintenance of cortical somatostatin interneuron subtype identity during migration* [Preprint]. Neuroscience. https://doi.org/10.1101/593285

Nawaratne, V., Leach, K., Suratman, N., Loiacono, R. E., Felder, C. C., Armbruster, B. N., Roth, B. L., Sexton, P. M., & Christopoulos, A. (2008). New Insights into the Function of M 4 Muscarinic Acetylcholine Receptors Gained Using a Novel Allosteric Modulator and a DREADD (Designer Receptor Exclusively Activated by a Designer Drug). Molecular Pharmacology, 74(4), 1119–1131. https://doi.org/10.1124/mol.108.049353

Neves, G., Shah, M. M., Liodis, P., Achimastou, A., Denaxa, M., Roalfe, G., Sesay, A., Walker, M. C., & Pachnis, V. (2013). The LIM Homeodomain Protein Lhx6 Regulates Maturation of Interneurons and Network Excitability in the Mammalian Cortex. Cerebral Cortex, 23(8), 1811–1823. https://doi.org/10.1093/cercor/bhs159

Newbern, J., Li, X., Shoemaker, S. E., Zhou, J., Zhong, J., Wu, Y., Bonder, D., Hollenback, S., Coppola, G., Geschwind, D. H., Landreth, G. E., & Snider, W. D. (2011). Specific functions for ERK/MAPK signaling during PNS development. Neuron, 69(1), 91–105. https://doi.org/10.1016/j.neuron.2010.12.003

Newbern, J., Zhong, J., Wickramasinghe, R. S., Li, X., Wu, Y., Samuels, I., Cherosky, N., Karlo, J. C., O’Loughlin, B., Wikenheiser, J., Gargesha, M., Doughman, Y. Q., Charron, J., Ginty, D. D., Watanabe, M., Saitta, S. C., Snider, W. D., & Landreth, G. E. (2008). Mouse and human phenotypes indicate a critical conserved role for ERK2 signaling in neural crest development. Proceedings of the National Academy of Sciences of the United States of America, 105(44), 17115–17120. https://doi.org/10.1073/pnas.0805239105

Nichols, J., Bjorklund, G. R., Newbern, J., & Anderson, T. (2018). Parvalbumin fast-spiking interneurons are selectively altered by paediatric traumatic brain injury. The Journal of Physiology, 596(7), 1277–1293. https://doi.org/10.1113/JP275393

Nigro, M. J., Hashikawa-Yamasaki, Y., & Rudy, B. (2018). Diversity and Connectivity of Layer 5 Somatostatin-Expressing Interneurons in the Mouse Barrel Cortex. The Journal of Neuroscience, 38(7), 1622–1633. https://doi.org/10.1523/JNEUROSCI.2415-17.2017

Nowaczyk, M. J. M., Thompson, B. A., Zeesman, S., Moog, U., Sanchez-Lara, P. A., Magoulas, P. L., Falk, R. E., Hoover-Fong, J. E., Batista, D. A. S., Amudhavalli, S. M., White, S. M., Graham, G. E., & Rauen, K. A. (2014). Deletion of MAP2K2/MEK2: A novel mechanism for a RASopathy? Clinical Genetics, 85(2), 138–146. https://doi.org/10.1111/cge.12116

Obrietan, K., Gao, X.-B., & van den Pol, A. N. (2002). Excitatory Actions of GABA Increase BDNF Expression via a MAPK-CREB–Dependent Mechanism—A Positive Feedback Circuit in Developing Neurons. Journal of Neurophysiology, 88(2), 1005–1015. https://doi.org/10.1152/jn.2002.88.2.1005

Oishi, K., Watatani, K., Itoh, Y., Okano, H., Guillemot, F., Nakajima, K., & Gotoh, Y. (2009). Selective induction of neocortical GABAergic neurons by the PDK1-Akt pathway through activation of Mash1. Proceedings of the National Academy of Sciences, 106(31), 13064–13069. https://doi.org/10.1073/pnas.0808400106

Okaty, B. W., Miller, M. N., Sugino, K., Hempel, C. M., & Nelson, S. B. (2009). Transcriptional and Electrophysiological Maturation of Neocortical Fast-Spiking GABAergic Interneurons. Journal of Neuroscience, 29(21), 7040–7052. https://doi.org/10.1523/JNEUROSCI.0105-09.2009

O’Leary, D. D. M., Chou, S.-J., & Sahara, S. (2007). Area Patterning of the Mammalian Cortex. Neuron, 56(2), 252–269. https://doi.org/10.1016/j.neuron.2007.10.010

Orduz, D., Benamer, N., Ortolani, D., Coppola, E., Vigier, L., Pierani, A., & Angulo, M. C. (2019). Developmental cell death regulates lineage-related interneuron-oligodendroglia functional clusters and oligodendrocyte homeostasis. Nature Communications, 10(1), 4249. https://doi.org/10.1038/s41467-019-11904-4

Ouwenga, R., Lake, A. M., Aryal, S., Lagunas, T., & Dougherty, J. D. (2018). The Differences in Local Translatome across Distinct Neuron Types Is Mediated by Both Baseline Cellular Differences and Post-transcriptional Mechanisms. Eneuro, 5(6), ENEURO.0320-18.2018. https://doi.org/10.1523/ENEURO.0320-18.2018

Owen, J. P., Chang, Y. S., Pojman, N. J., Bukshpun, P., Wakahiro, M. L. J., Marco, E. J., Berman, J. I., Spiro, J. E., Chung, W. K., Buckner, R. L., Roberts, T. P. L., Nagarajan, S. S., Sherr, E. H., Mukherjee, P., & Simons VIP Consortium. (2014). Aberrant white matter microstructure in children with 16p11.2 deletions. The Journal of Neuroscience: The Official Journal of the Society for Neuroscience, 34(18), 6214–6223. https://doi.org/10.1523/JNEUROSCI.4495-13.2014

Owens, D. F., & Kriegstein, A. R. (2002). Is there more to gaba than synaptic inhibition? Nature Reviews Neuroscience, 3(9), 715–727. https://doi.org/10.1038/nrn919

Pan, N. C., Fang, A., Shen, C., Sun, L., Wu, Q., & Wang, X. (2018). Early Excitatory Activity-Dependent Maturation of Somatostatin Interneurons in Cortical Layer 2/3 of Mice. Cerebral Cortex. https://doi.org/10.1093/cercor/bhy293

Pantazopoulos, H., Wiseman, J. T., Markota, M., Ehrenfeld, L., & Berretta, S. (2017). Decreased Numbers of Somatostatin-Expressing Neurons in the Amygdala of Subjects With Bipolar Disorder or Schizophrenia: Relationship to Circadian Rhythms. Biological Psychiatry, 81(6), 536–547. https://doi.org/10.1016/j.biopsych.2016.04.006

Paul, A., Crow, M., Raudales, R., He, M., Gillis, J., & Huang, Z. J. (2017). Transcriptional Architecture of Synaptic Communication Delineates GABAergic Neuron Identity. Cell, 171(3), 522–539.e20. https://doi.org/10.1016/j.cell.2017.08.032

Payne, J. M., Hearps, S. J. C., Walsh, K. S., Paltin, I., Barton, B., Ullrich, N. J., Haebich, K. M., Coghill, D., Gioia, G. A., Cantor, A., Cutter, G., Tonsgard, J. H., Viskochil, D., Rey-Casserly, C., Schorry, E. K., Ackerson, J. D., Klesse, L., Fisher, M. J., Gutmann, D. H., … the NF Clinical Trials Consortium. (2019). Reproducibility of cognitive endpoints in clinical trials: Lessons from neurofibromatosis type 1. Annals of Clinical and Translational Neurology, 6(12), 2555–2565. https://doi.org/10.1002/acn3.50952

Peerboom, C., & Wierenga, C. J. (2021). The postnatal GABA shift: A developmental perspective. Neuroscience & Biobehavioral Reviews, 124, 179–192. https://doi.org/10.1016/j.neubiorev.2021.01.024

Peng, Z., Zhang, N., Wei, W., Huang, C. S., Cetina, Y., Otis, T. S., & Houser, C. R. (2013). A Reorganized GABAergic Circuit in a Model of Epilepsy: Evidence from Optogenetic Labeling and Stimulation of Somatostatin Interneurons. Journal of Neuroscience, 33(36), 14392–14405. https://doi.org/10.1523/JNEUROSCI.2045-13.2013

Petros, T. J., Bultje, R. S., Ross, M. E., Fishell, G., & Anderson, S. A. (2015). Apical versus Basal Neurogenesis Directs Cortical Interneuron Subclass Fate. Cell Reports, 13(6), 1090–1095. https://doi.org/10.1016/j.celrep.2015.09.079

Pietrobono, S., Gagliardi, S., & Stecca, B. (2019). Non-canonical Hedgehog Signaling Pathway in Cancer: Activation of GLI Transcription Factors Beyond Smoothened. Frontiers in Genetics, 10, 556. https://doi.org/10.3389/fgene.2019.00556

Po, A., Silvano, M., Miele, E., Capalbo, C., Eramo, A., Salvati, V., Todaro, M., Besharat, Z. M., Catanzaro, G., Cucchi, D., Coni, S., Di Marcotullio, L., Canettieri, G., Vacca, A., Stassi, G., De Smaele, E., Tartaglia, M., Screpanti, I., De Maria, R., & Ferretti, E. (2017). Noncanonical GLI1 signaling promotes stemness features and in vivo growth in lung adenocarcinoma. Oncogene, 36(32), 4641–4652. https://doi.org/10.1038/onc.2017.91

Polleux, F., Whitford, K. L., Dijkhuizen, P. A., Vitalis, T., & Ghosh, A. (2002). Control of cortical interneuron migration by neurotrophins and PI3-kinase signaling. *Development (Cambridge*, England), 129(13), 3147–3160.

Priya, R., Paredes, M. F., Karayannis, T., Yusuf, N., Liu, X., Jaglin, X., Graef, I., Alvarez-Buylla, A., & Fishell, G. (2018). Activity Regulates Cell Death within Cortical Interneurons through a Calcineurin-Dependent Mechanism. Cell Reports, 22(7), 1695–1709. https://doi.org/10.1016/j.celrep.2018.01.007

Pucilowska, J., Puzerey, P. A., Karlo, J. C., Galan, R. F., & Landreth, G. E. (2012). Disrupted ERK Signaling during Cortical Development Leads to Abnormal Progenitor Proliferation, Neuronal and Network Excitability and Behavior, Modeling Human Neuro-Cardio-Facial-Cutaneous and Related Syndromes. Journal of Neuroscience, 32(25), 8663–8677. https://doi.org/10.1523/JNEUROSCI.1107-12.2012

Pucilowska, J., Vithayathil, J., Tavares, E. J., Kelly, C., Karlo, J. C., & Landreth, G. E. (2015). The 16p11.2 Deletion Mouse Model of Autism Exhibits Altered Cortical Progenitor Proliferation and Brain Cytoarchitecture Linked to the ERK MAPK Pathway. Journal of Neuroscience, 35(7), 3190–3200. https://doi.org/10.1523/JNEUROSCI.4864-13.2015

Qureshi, A. Y., Mueller, S., Snyder, A. Z., Mukherjee, P., Berman, J. I., Roberts, T. P. L., Nagarajan, S. S., Spiro, J. E., Chung, W. K., Sherr, E. H., Buckner, R. L., & Simons VIP Consortium. (2014). Opposing brain differences in 16p11.2 deletion and duplication carriers. The Journal of Neuroscience: The Official Journal of the Society for Neuroscience, 34(34), 11199–11211. https://doi.org/10.1523/JNEUROSCI.1366-14.2014

Rauen, K. A. (2013). The RASopathies. Annual Review of Genomics and Human Genetics, 14(1), 355–369. https://doi.org/10.1146/annurev-genom-091212-153523

Reh, R. K., Dias, B. G., Nelson, C. A., Kaufer, D., Werker, J. F., Kolb, B., Levine, J. D., & Hensch, T. K. (2020). Critical period regulation across multiple timescales. Proceedings of the National Academy of Sciences, 201820836. https://doi.org/10.1073/pnas.1820836117

Rizvi, T. A., Akunuru, S., de Courten-Myers, G., Switzer, R. C., Nordlund, M. L., & Ratner, N. (1999). Region-specific astrogliosis in brains of mice heterozygous for mutations in the neurofibromatosis type 1 (Nf1) tumor suppressor. Brain Research, 816(1), 111–123. https://doi.org/10.1016/s0006-8993(98)01133-0

Robbins, R. J., Brines, M. L., Kim, J. H., Adrian, T., De Lanerolle, N., Welsh, S., & Spencer, D. D. (1991). A selective loss of somatostatin in the hippocampus of patients with temporal lobe epilepsy. Annals of Neurology, 29(3), 325–332. https://doi.org/10.1002/ana.410290316

Rogan, S. C., & Roth, B. L. (2011). Remote Control of Neuronal Signaling. Pharmacological Reviews, 63(2), 291–315. https://doi.org/10.1124/pr.110.003020

Rogers, S., Rozman, P. A., Valero, M., Doyle, W. K., & Buzsáki, G. (2021). Mechanisms and plasticity of chemogenically induced interneuronal suppression of principal cells. Proceedings of the National Academy of Sciences, 118(2), e2014157118. https://doi.org/10.1073/pnas.2014157118

Rosina, E., Battan, B., Siracusano, M., Di Criscio, L., Hollis, F., Pacini, L., Curatolo, P., & Bagni, C. (2019). Disruption of mTOR and MAPK pathways correlates with severity in idiopathic autism. Translational Psychiatry, 9(1), 50. https://doi.org/10.1038/s41398-018-0335-z

Roth, B. L. (2016). DREADDs for Neuroscientists. Neuron, 89(4), 683–694. https://doi.org/10.1016/j.neuron.2016.01.040

Ryu, H.-H., Kang, M., Park, J., Park, S.-H., & Lee, Y.-S. (2019). Enriched expression of NF1 in inhibitory neurons in both mouse and human brain. Molecular Brain, 12(1), 60. https://doi.org/10.1186/s13041-019-0481-0

Ryu, H.-H., Kim, T., Kim, J.-W., Kang, M., Park, P., Kim, Y. G., Kim, H., Ha, J., Choi, J. E., Lee, J., Lim, C.-S., Kim, C.-H., Kim, S. J., Silva, A. J., Kaang, B.-K., & Lee, Y.-S. (2019). Excitatory neuron–specific SHP2-ERK signaling network regulates synaptic plasticity and memory. Science Signaling, 12(571), eaau5755. https://doi.org/10.1126/scisignal.aau5755

Saitta, S. C., Harris, S. E., McDonald-McGinn, D. M., Emanuel, B. S., Tonnesen, M. K., Zackai, E. H., Seitz, S. C., & Driscoll, D. A. (2004). Independent de novo 22q11.2 deletions in first cousins with DiGeorge/velocardiofacial syndrome. American Journal of Medical Genetics, 124A(3), 313–317. https://doi.org/10.1002/ajmg.a.20421

Samuels, I. S., Karlo, J. C., Faruzzi, A. N., Pickering, K., Herrup, K., Sweatt, J. D., Saitta, S. C., & Landreth, G. E. (2008). Deletion of ERK2 Mitogen-Activated Protein Kinase Identifies Its Key Roles in Cortical Neurogenesis and Cognitive Function. Journal of Neuroscience, 28(27), 6983– 6995. https://doi.org/10.1523/JNEUROSCI.0679-08.2008

Samuels, I. S., Saitta, S. C., & Landreth, G. E. (2009). MAP’ing CNS Development and Cognition: An ERKsome Process. Neuron, 61(2), 160–167. https://doi.org/10.1016/j.neuron.2009.01.001

Sánchez, A. I., García-Acero, M. A., Paredes, A., Quero, R., Ortega, R. I., Rojas, J. A., Herrera, D., Parra, M., Prieto, K., Ángel, J., Rodríguez, L.-S., Prieto, J. C., & Franco, M. (2020). Immunodeficiency in a Patient with 22q11.2 Distal Deletion Syndrome and a p.Ala7dup Variant in the **MAPK1** Gene. *Molecular Syndromology*, *11*(1), 15–23. https://doi.org/10.1159/000506032

Sanchez-Ortiz, E., Cho, W., Nazarenko, I., Mo, W., Chen, J., & Parada, L. F. (2014). NF1 regulation of RAS/ERK signaling is required for appropriate granule neuron progenitor expansion and migration in cerebellar development. Genes & Development, 28(21), 2407–2420. https://doi.org/10.1101/gad.246603.114

Sanz, E., Yang, L., Su, T., Morris, D. R., McKnight, G. S., & Amieux, P. S. (2009). Cell-type-specific isolation of ribosome-associated mRNA from complex tissues. Proceedings of the National Academy of Sciences, 106(33), 13939–13944. https://doi.org/10.1073/pnas.0907143106

Selcher, J. C. (2001). Mice Lacking the ERK1 Isoform of MAP Kinase Are Unimpaired in Emotional Learning. Learning & Memory, 8(1), 11–19. https://doi.org/10.1101/lm.37001

Shimamura, K., & Rubenstein, J. L. (1997). Inductive interactions direct early regionalization of the mouse forebrain. Development, 124(14), 2709–2718. https://doi.org/10.1242/dev.124.14.2709

Sik, A., Hajos, N., Gulacsi, A., Mody, I., & Freund, T. F. (1998). The absence of a major Ca2+ signaling pathway in GABAergic neurons of the hippocampus. Proceedings of the National Academy of Sciences, 95(6), 3245–3250. https://doi.org/10.1073/pnas.95.6.3245

Silva, A. I., Ehrhart, F., Ulfarsson, M. O., Stefansson, H., Stefansson, K., Wilkinson, L. S., Hall, J., & Linden, D. E. J. (2022). Neuroimaging Findings in Neurodevelopmental Copy Number Variants: Identifying Molecular Pathways to Convergent Phenotypes. Biological Psychiatry, 92(5), 341–361. https://doi.org/10.1016/j.biopsych.2022.03.018

Smith, S. J., Sümbül, U., Graybuck, L. T., Collman, F., Seshamani, S., Gala, R., Gliko, O., Elabbady, L., Miller, J. A., Bakken, T. E., Rossier, J., Yao, Z., Lein, E., Zeng, H., Tasic, B., & Hawrylycz, M. (2019). Single-cell transcriptomic evidence for dense intracortical neuropeptide networks. ELife, 8, e47889. https://doi.org/10.7554/eLife.47889

Song, Y.-H., Hwang, Y.-S., Kim, K., Lee, H.-R., Kim, J.-H., Maclachlan, C., Dubois, A., Jung, M. W., Petersen, C. C. H., Knott, G., Lee, S.-H., & Lee, S.-H. (2020). Somatostatin enhances visual processing and perception by suppressing excitatory inputs to parvalbumin-positive interneurons in V1. Science Advances, 6(17), eaaz0517. https://doi.org/10.1126/sciadv.aaz0517

Song, Y.-H., Yoon, J., & Lee, S.-H. (2021). The role of neuropeptide somatostatin in the brain and its application in treating neurological disorders. Experimental & Molecular Medicine, 53(3), 328–338. https://doi.org/10.1038/s12276-021-00580-4

Southwell, D. G., Paredes, M. F., Galvao, R. P., Jones, D. L., Froemke, R. C., Sebe, J. Y., Alfaro-Cervello, C., Tang, Y., Garcia-Verdugo, J. M., Rubenstein, J. L., Baraban, S. C., & Alvarez-Buylla, A. (2012). Intrinsically determined cell death of developing cortical interneurons. Nature, 491(7422), 109–113. https://doi.org/10.1038/nature11523

Spiegel, I., Mardinly, A. R., Gabel, H. W., Bazinet, J. E., Couch, C. H., Tzeng, C. P., Harmin, D. A., & Greenberg, M. E. (2014). Npas4 Regulates Excitatory-Inhibitory Balance within Neural Circuits through Cell-Type-Specific Gene Programs. Cell, 157(5), 1216–1229. https://doi.org/10.1016/j.cell.2014.03.058

Sun, Y., Qian, L., Xu, L., Hunt, S., & Sah, P. (2020). Somatostatin neurons in the central amygdala mediate anxiety by disinhibition of the central sublenticular extended amygdala. Molecular Psychiatry. https://doi.org/10.1038/s41380-020-00894-1

Tallent, M. K., & Qiu, C. (2008). Somatostatin: An endogenous antiepileptic. Molecular and Cellular Endocrinology, 286(1–2), 96–103. https://doi.org/10.1016/j.mce.2007.12.004

Talley, M. J., Nardini, D., Qin, S., Prada, C. E., Ehrman, L. A., & Waclaw, R. R. (2021). A role for sustained MAPK activity in the mouse ventral telencephalon. Developmental Biology, 476, 137–147. https://doi.org/10.1016/j.ydbio.2021.03.019

Taniguchi, H., He, M., Wu, P., Kim, S., Paik, R., Sugino, K., Kvitsani, D., Fu, Y., Lu, J., Lin, Y., Miyoshi, G., Shima, Y., Fishell, G., Nelson, S. B., & Huang, Z. J. (2011). A Resource of Cre Driver Lines for Genetic Targeting of GABAergic Neurons in Cerebral Cortex. Neuron, 71(6), 995–1013. https://doi.org/10.1016/j.neuron.2011.07.026

Tasic, B., Menon, V., Nguyen, T. N., Kim, T. K., Jarsky, T., Yao, Z., Levi, B., Gray, L. T., Sorensen, S. A., Dolbeare, T., Bertagnolli, D., Goldy, J., Shapovalova, N., Parry, S., Lee, C., Smith, K., Bernard, A., Madisen, L., Sunkin, S. M., … Zeng, H. (2016). Adult mouse cortical cell taxonomy revealed by single cell transcriptomics. Nature Neuroscience, 19(2), 335–346. https://doi.org/10.1038/nn.4216

Tereshko, L., Gao, Y., Cary, B. A., Turrigiano, G. G., & Sengupta, P. (2021). Ciliary neuropeptidergic signaling dynamically regulates excitatory synapses in postnatal neocortical pyramidal neurons. ELife, 10, e65427. https://doi.org/10.7554/eLife.65427

Thomas, G. M., & Huganir, R. L. (2004). MAPK cascade signalling and synaptic plasticity. Nature Reviews Neuroscience, 5(3), 173–183. https://doi.org/10.1038/nrn1346

Tidyman, W. E., & Rauen, K. A. (2016). Pathogenetics of the RASopathies. Human Molecular Genetics, 25(R2), R123–R132. https://doi.org/10.1093/hmg/ddw191

Ting, A. K., Chen, Y., Wen, L., Yin, D.-M., Shen, C., Tao, Y., Liu, X., Xiong, W.-C., & Mei, L. (2011). Neuregulin 1 Promotes Excitatory Synapse Development and Function in GABAergic Interneurons. Journal of Neuroscience, 31(1), 15–25. https://doi.org/10.1523/JNEUROSCI.2538-10.2011

Topolnik, L. (2012). Dendritic calcium mechanisms and long-term potentiation in cortical inhibitory interneurons: Calcium in interneuron dendrites. European Journal of Neuroscience, 35(4), 496–506. https://doi.org/10.1111/j.1460-9568.2011.07988.x

Topolnik, L., Azzi, M., Morin, F., Kougioumoutzakis, A., & Lacaille, J.-C. (2006). mGluR1/5 subtype-specific calcium signalling and induction of long-term potentiation in rat hippocampal oriens/alveus interneurones: MGluR1/5 Ca ^2+^ signalling and LTP in interneurones. The Journal of Physiology, 575(1), 115–131. https://doi.org/10.1113/jphysiol.2006.112896

Tripp, A., Kota, R. S., Lewis, D. A., & Sibille, E. (2011). Reduced somatostatin in subgenual anterior cingulate cortex in major depression. Neurobiology of Disease, 42(1), 116–124. https://doi.org/10.1016/j.nbd.2011.01.014

Tucker, E. S., Segall, S., Gopalakrishna, D., Wu, Y., Vernon, M., Polleux, F., & LaMantia, A.-S. (2008). Molecular Specification and Patterning of Progenitor Cells in the Lateral and Medial Ganglionic Eminences. Journal of Neuroscience, 28(38), 9504–9518. https://doi.org/10.1523/JNEUROSCI.2341-08.2008

Tuncdemir, S. N., Wamsley, B., Stam, F. J., Osakada, F., Goulding, M., Callaway, E. M., Rudy, B., & Fishell, G. (2016). Early Somatostatin Interneuron Connectivity Mediates the Maturation of Deep Layer Cortical Circuits. Neuron, 89(3), 521–535. https://doi.org/10.1016/j.neuron.2015.11.020

Tyson, J. A., Goldberg, E. M., Maroof, A. M., Xu, Q., Petros, T. J., & Anderson, S. A. (2015). Duration of culture and sonic hedgehog signaling differentially specify PV versus SST cortical interneuron fates from embryonic stem cells. Development, 142(7), 1267–1278. https://doi.org/10.1242/dev.111526

Urban-Ciecko, J., & Barth, A. L. (2016). Somatostatin-expressing neurons in cortical networks. Nature Reviews Neuroscience, 17(7), 401–409. https://doi.org/10.1038/nrn.2016.53

Vithayathil, J., Pucilowska, J., Goodnough, L. H., Atit, R. P., & Landreth, G. E. (2015). Dentate Gyrus Development Requires ERK Activity to Maintain Progenitor Population and MAPK Pathway Feedback Regulation. Journal of Neuroscience, 35(17), 6836–6848. https://doi.org/10.1523/JNEUROSCI.4196-14.2015

Vogt, D., Cho, K. K. A., Lee, A. T., Sohal, V. S., & Rubenstein, J. L. R. (2015). The Parvalbumin/Somatostatin Ratio Is Increased in Pten Mutant Mice and by Human PTEN ASD Alleles. Cell Reports, 11(6), 944–956. https://doi.org/10.1016/j.celrep.2015.04.019

Vong, L., Ye, C., Yang, Z., Choi, B., Chua, S., & Lowell, B. B. (2011). Leptin Action on GABAergic Neurons Prevents Obesity and Reduces Inhibitory Tone to POMC Neurons. Neuron, 71(1), 142–154. https://doi.org/10.1016/j.neuron.2011.05.028

Wei, Y., Han, X., & Zhao, C. (2020). PDK1 regulates the survival of the developing cortical interneurons. Molecular Brain, 13(1), 65. https://doi.org/10.1186/s13041-020-00604-6

Wengert, E. R., Miralles, R. M., Wedgwood, K. C. A., Wagley, P. K., Strohm, S. M., Panchal, P. S., Idrissi, A. M., Wenker, I. C., Thompson, J. A., Gaykema, R. P., & Patel, M. K. (2021). Somatostatin-Positive Interneurons Contribute to Seizures in SCN8A Epileptic Encephalopathy. The Journal of Neuroscience, 41(44), 9257–9273. https://doi.org/10.1523/JNEUROSCI.0718-21.2021

Wonders, C. P., & Anderson, S. A. (2006). The origin and specification of cortical interneurons. Nature Reviews Neuroscience, 7(9), 687–696. https://doi.org/10.1038/nrn1954

Wong, F. K., Bercsenyi, K., Sreenivasan, V., Portalés, A., Fernández-Otero, M., & Marín, O. (2018). Pyramidal cell regulation of interneuron survival sculpts cortical networks. Nature, 557(7707), 668–673. https://doi.org/10.1038/s41586-018-0139-6

Wu, G.-Y., Deisseroth, K., & Tsien, R. W. (2001). Activity-dependent CREB phosphorylation: Convergence of a fast, sensitive calmodulin kinase pathway and a slow, less sensitive mitogen-activated protein kinase pathway. Proceedings of the National Academy of Sciences, 98(5), 2808– 2813. https://doi.org/10.1073/pnas.051634198

Xing, J., Ginty, D. D., & Greenberg, M. E. (1996). Coupling of the RAS-MAPK Pathway to Gene Activation by RSK2, a Growth Factor-Regulated CREB Kinase. Science, 273(5277), 959–963. https://doi.org/10.1126/science.273.5277.959

Xing, L., Larsen, R. S., Bjorklund, G. R., Li, X., Wu, Y., Philpot, B. D., Snider, W. D., & Newbern, J. M. (2016). Layer specific and general requirements for ERK/MAPK signaling in the developing neocortex. ELife, 5, e11123. https://doi.org/10.7554/eLife.11123

Xu, Q., Guo, L., Moore, H., Waclaw, R. R., Campbell, K., & Anderson, S. A. (2010). Sonic Hedgehog Signaling Confers Ventral Telencephalic Progenitors with Distinct Cortical Interneuron Fates. Neuron, 65(3), 328–340. https://doi.org/10.1016/j.neuron.2010.01.004

Xu, Q., Tam, M., & Anderson, S. A. (2008). Fate mapping Nkx2.1-lineage cells in the mouse telencephalon. The Journal of Comparative Neurology, 506(1), 16–29. https://doi.org/10.1002/cne.21529

Xu, Q., Wonders, C. P., & Anderson, S. A. (2005). Sonic hedgehog maintains the identity of cortical interneuron progenitors in the ventral telencephalon. Development, 132(22), 4987–4998. https://doi.org/10.1242/dev.02090

Yamamoto, K. K., Gonzalez, G. A., Biggs, W. H., & Montminy, M. R. (1988). Phosphorylation-induced binding and transcriptional efficacy of nuclear factor CREB. Nature, 334(6182), 494–498. https://doi.org/10.1038/334494a0

Yap, E.-L., & Greenberg, M. E. (2018). Activity-Regulated Transcription: Bridging the Gap between Neural Activity and Behavior. Neuron, 100(2), 330–348. https://doi.org/10.1016/j.neuron.2018.10.013

Yu, W., Wang, Y., McDonnell, K., Stephen, D., & Bai, C. B. (2009). Patterning of ventral telencephalon requires positive function of Gli transcription factors. Developmental Biology, 334(1), 264–275. https://doi.org/10.1016/j.ydbio.2009.07.026

Zeyda, T., Diehl, N., Paylor, R., Brennan, M. B., & Hochgeschwender, U. (2001). Impairment in motor learning of somatostatin null mutant mice. Brain Research, 906(1–2), 107–114. https://doi.org/10.1016/S0006-8993(01)02563-X

Zhu, H., Aryal, D. K., Olsen, R. H. J., Urban, D. J., Swearingen, A., Forbes, S., Roth, B. L., & Hochgeschwender, U. (2016). Cre-dependent DREADD (Designer Receptors Exclusively Activated by Designer Drugs) mice. *Genesis (New York*, N.Y.: 2000), *54*(8), 439–446. https://doi.org/10.1002/dvg.22949

Zou, D., Chen, L., Deng, D., Jiang, D., Dong, F., McSweeney, C., Zhou, Y., Liu, L., Chen, G., Wu, Y., & Mao, Y. (2016). DREADD in Parvalbumin Interneurons of the Dentate Gyrus Modulates Anxiety, Social Interaction and Memory Extinction. Current Molecular Medicine, 16(1), 91–102. https://doi.org/10.2174/1566524016666151222150024

